# Tankyrase inhibition demonstrates anti-fibrotic effects in preclinical pulmonary fibrosis models

**DOI:** 10.1101/2025.11.13.688191

**Authors:** Shoshy Alam Brinch, Ida Johnsen, Lena Willmer, Ole Vidhammer Bjornstad, Teresa Lucifora, Karl Martin Forbord, Maria Candamo-Lourido, Maureen Tania Meling, Ingrid Lofthus Morken, Martin Frank Strand, Danny Jonigk, Patrick Zardo, Hans Gerd Fieguth, Christina Hesse, Mark Alexander Skarsfeldt, Morten Asser Karsdal, Erika Ferrari, Paola Occhetta, Roberta Visone, Stephen Jordan, Joseph Lee, Harry Holton, Simon Rayner, Anita Wegert, Simon Cruwys, Stefan Krauss, Jo Waaler

## Abstract

**Background:** Idiopathic pulmonary fibrosis (IPF) is a progressive and fatal lung disease with limited treatment options. Although transforming growth factor beta 1 (TGFB1, TGFβ) is a key driver of fibrosis, additional signaling pathways, including wingless-type mammary tumor virus integration site (WNT)/β-catenin and yes-associated protein 1 (YAP), contribute to IPF pathogenesis. Clinical data indicate that inhibition of TGFβ alone provides limited efficacy or is associated with toxicity, underscoring the need for alternative therapeutic approaches. Tankyrase (TNKS) 1 and 2 are post-translational regulators of WNT/β-catenin and YAP signaling and therefore represent promising antifibrotic targets. OM-153, a potent and selective TNKS inhibitor, exhibits pharmacological properties suitable for preclinical development in IPF.

**Methods:** Primary normal human lung fibroblasts (NHLF), Scar-in-a-Jar assays, lung-on-a-chip models, and precision-cut lung slices (PCLS) from non-pulmonary fibrosis (non-PF) tissue were stimulated with an IPF-relevant cytokine cocktail (IPF-RC) designed to accurately recapitulate the pro-fibrotic environment and compared to TGFβ. These models, with bleomycin-challenged mice and PCLS from end-stage pulmonary fibrosis (PF) patients, were treated with OM-153. Fibrosis markers, extracellular matrix (ECM) components, and signaling pathway-specific gene expression or protein markers were assessed by real-time qRT-PCR, RNA sequencing, immunoblotting, ELISA, and immunofluorescence.

**Results:** OM-153 stabilized the direct TNKS targets axin 1 (AXIN1) and angiomotin-like 1 (AMOTL1), suppressed WNT/β-catenin and YAP signaling. In parallel, it reduced profibrotic ECM expression across *in vitro*, *in vivo*, and *ex vivo* IPF models.

**Conclusions:** Selective TNKS inhibition by OM-153 demonstrates broad antifibrotic activity in multiple preclinical models, supporting further development as a potential disease-modifying strategy for IPF.

**Shareable abstract:** Our findings show that the potent and selective TNKS inhibitor OM-153 suppresses WNT/β-catenin and YAP signaling, reducing pro-fibrotic ECM expression in preclinical IPF models, supporting TNKS inhibition as a novel antifibrotic strategy.

## Introduction

Interstitial lung disease (ILD) comprises a heterogeneous group of pulmonary disorders arising from autoimmune conditions, infectious agents, and genetic abnormalities [1, 2]. These disorders are characterized by progressive scarring of the lung parenchyma, leading to increased stiffness, impaired gas exchange, and ultimately respiratory failure [2]. IPF is the most severe ILD subtype, with unknown etiology, a median survival of ∼3 years post-diagnosis, and an incidence of 3–9 cases per 100,000 adults [3, 4]. Current standard-of-care (SoC) therapies, pirfenidone and nintedanib, modestly slow disease progression but neither halt or reverse fibrosis and are associated with significant adverse effects [2]. Over the past decade, drug development has predominantly targeted the TGFβ signaling pathway, however, approaches including olitigaltin, bexotegrast, and other pathway-directed agents have uniformly failed in clinical trials due to insufficient efficacy or unacceptable toxicity [4, 5]. This highlights the need to investigate non–TGFβ-directed therapies capable of effectively slowing or reversing fibrotic progression in IPF.

Alongside TGFβ, WNT/β-catenin and YAP signaling represent central pathways in lung fibrosis [6]. Dysregulated activation of these pathways promotes epithelial–mesenchymal transition (EMT), fibroblast-to-myofibroblast transition (FMT), fibroblast proliferation, ECM deposition, and airway remodeling, thereby contributing to IPF pathogenesis [7–9]. Pharmacological inhibition of WNT/β-catenin and YAP signaling attenuates fibrosis in preclinical models, underscoring their therapeutic potential [10, 11]. Despite this, no WNT/β-catenin- or YAP-targeting therapies are clinically approved for IPF or any other disease.

Traditionally, fibrogenesis in fibroblasts has been induced by TGFβ stimulation alone [4]. This approach, however, may preferentially identify compounds targeting TGFβ signaling and is less informative for evaluating other mechanisms, including TNKS inhibition by OM-153, which exerts downstream effects on WNT/β-catenin and YAP signaling [4]. The IPF-relevant cytokine cocktail (IPF-RC), comprising nine cytokines, was recently established to more accurately reproduce the pro-fibrotic milieu of human IPF lungs, activating multiple signaling pathways that collectively drive fibrogenesis [12].

The biotargets TNKS 1 and 2 are multifunctional regulators implicated in several signaling pathways and represent key nodes for modulating WNT/β-catein and YAP activity [13]. Mechanistically, TNKS catalyzes the poly(ADP-ribosyl)ation of axin (AXIN) and angiomotin (AMOT) proteins, targeting them for proteasomal degradation via the ubiquitin–proteasome system [13–16]. TNKS inhibition stabilizes AXINs, the rate-limiting scaffold of the β-catenin destruction complex, thereby enhancing β-catenin degradation and suppressing WNT/β-catenin signaling [17–19]. Similarly, TNKS inhibition stabilizes AMOT proteins, which sequester YAP and WW domain-containing transcription regulator 1 (WWTR1/TAZ) in the cytoplasm, attenuating YAP signaling [16, 18]. Proof-of-concept studies demonstrated that early-generation TNKS inhibitors reduced fibrosis in skin fibroblasts and in bleomycin-induced dermal and pulmonary fibrosis models in mice [20–22]. More recently, AstraZeneca’s porcupine and WNT inhibitor AZD5055 showed efficacy in preclinical fibrosis models and favorable safety and tolerability in a phase 1 study [23].

Significant efforts have been directed toward the development of highly selective small-molecule TNKS inhibitors [19, 24–27]. Clinical translation, however, has been halted by concerns regarding on-target and pathway-specific toxicities, notably intestinal toxicity [28–30]. Rational design strategies targeting the adenosine-binding pocket of TNKS catalytic domains, including 1,2,4-triazole–based scaffolds, have yielded inhibitors with improved potency and selectivity [31–33].

Among these, OM-153 is a next-generation preclinical TNKS inhibitor that demonstrates picomolar IC₅₀ activity in cellular WNT/β-catenin reporter assays and shows no detectable off-target liabilities [33, 34]. OM-153 also exhibits favorable absorption, distribution, metabolism, and excretion (ADME) properties, along with an optimized peroral pharmacokinetic profile providing a 30-fold therapeutic window in mice [33, 34].

Parallel progress has also advanced the selective TNKS inhibitor basroparib (STP1002) and the non-selective TNKS inhibitor stenoparib (2X-121), both of which have shown safety and tolerability in phase 1 clinical oncology studies [35, 36].

Here, we demonstrate that the next-generation TNKS inhibitor OM-153 stabilizes AXIN and AMOT proteins, thereby suppressing WNT/β-catenin and YAP signaling. Across diverse preclinical models of lung fibrosis, OM-153 robustly attenuates ECM synthesis and deposition. Collectively, these findings identify TNKS inhibition as a mechanistically distinct and promising therapeutic approach for IPF.

## Materials and Methods

Detailed methods are provided in the supplementary material.

### Human lung fibroblast assays

NHLFs were stimulated with IPF-RC or TGFβ [12] and treated with small-molecule inhibitors according to established protocols [18]. Immunoblotting, RNA isolation, and real-time qRT-PCR analyses were performed as previously described [18].

### Mouse bleomycin-induced lung fibrosis model

Animal experiments were conducted at WuXi AppTec (Shanghai, China) following their standard protocol, approved by the local authorities, and in accordance with FELASA guidelines and institutional recommendations.

### Human precision-cut lung slices

PCLS from non-PF or end-stage PF donor tissue were stimulated with IPF-RC and treated with small-molecule inhibitors.

### Statistical analysis

Statistical analyses were performed as previously described using t-tests and Mann-Whitney rank sum tests [18].

## Results

### OM-153 reduces IPF-RC-stimulated ECM gene and protein expression in normal human lung fibroblasts

To compare ECM induction by IPF-RC versus TGFβ, NHLFs were stimulated for 72 hours in a 2D *in vitro* model. The expression of the clinically relevant collagen markers *COL3A1* and *COL6A1*, which are associated with IPF progression [37], as well as *COL1A1*, was assessed. Concentration–response analysis revealed that IPF-RC (0.1–10×) significantly upregulated *COL1A1*, *COL3A1*, and *COL6A1*, whereas TGFβ stimulation alone (0.1–10 ng/mL) induced only *COL1A1* and *COL3A1* expression (**supplementary figure 1**). The anti-fibrotic efficacy of OM-153 was then evaluated in NHLFs stimulated side-by-side with either IPF-RC (1X, including 0.3 ng/mL TGFβ) or TGFβ (0.3 ng/mL) (**figure 1a**,**b**). Real-time qRT-PCR analysis showed that IPF-RC stimulation increased *COL1A1* (2.1-fold), *COL3A1* (3.0-fold), and *COL6A1* (3.4-fold) expression relative to the control (**figure 1a**). OM-153 significantly reduced all three ECM markers in a concentration-dependent manner, whereas nintedanib only moderately suppressed their expression. In contrast, TGFβ stimulation increased only *COL1A1* (1.7-fold) and *COL3A1* (1.3-fold), while the *COL6A1* remained unchanged (0.9-fold), and neither OM-153 nor nintedanib affected these markers (**figure 1b**). These results demonstrate that IPF-RC more accurately recapitulates the pro-fibrotic environment of human IPF compared with TGFβ alone, and that the TNKS inhibitor OM-153 potently inhibits ECM marker upregulation in this context.

**Figure 1.**
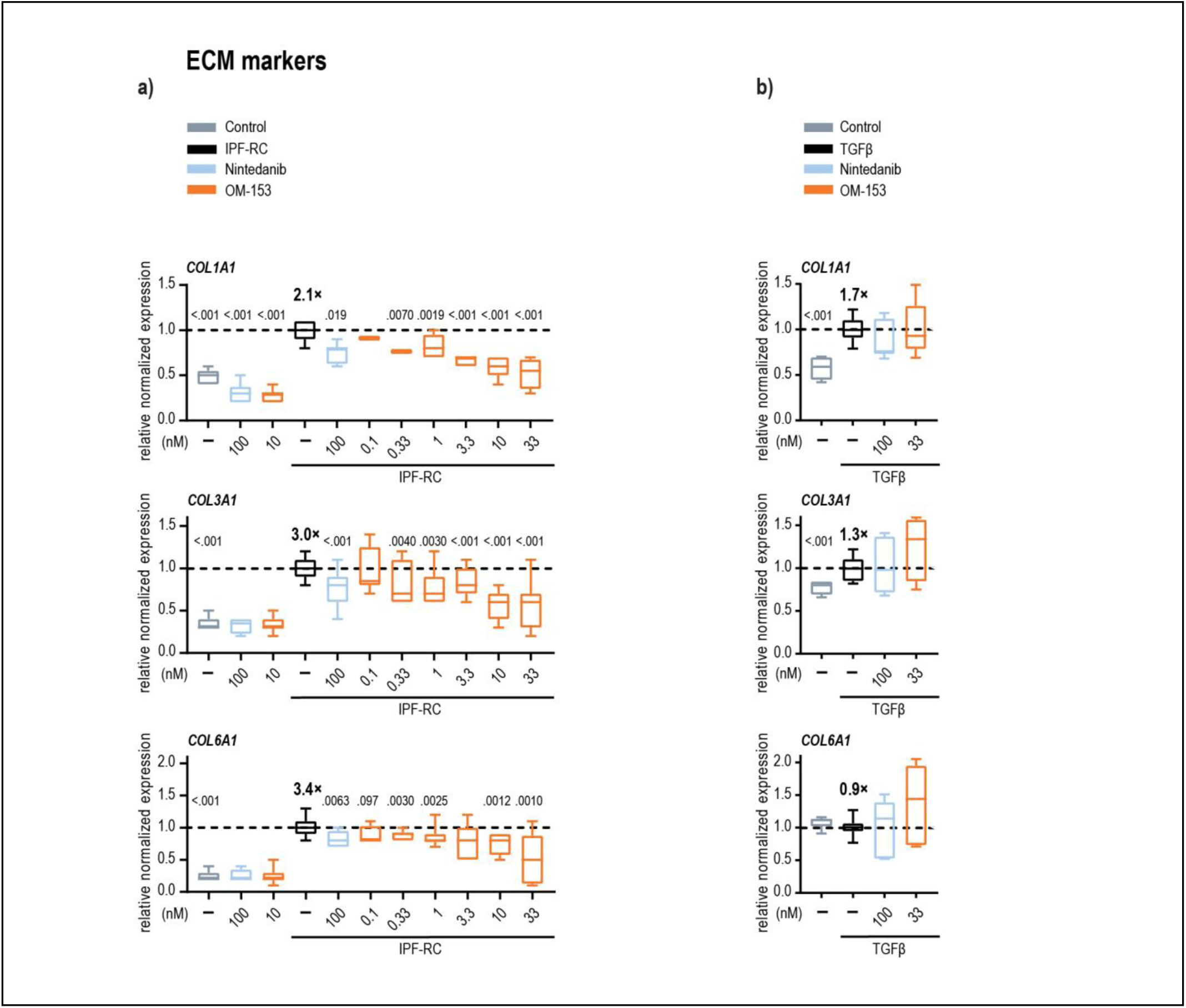
OM-153 reduces IPF-RC-stimulated ECM gene expression in normal human lung fibroblasts a, **b)** Real-time qRT-PCR of ECM markers *COL1A1*, *COL3A1*, and *COL6A1* in NHLFs cultured in 2D for 72 hours. Cells were stimulated with **a)** IPF-RC (1×) or **b)** TGFβ (0.3 ng/mL) (both, black), and treated with nintedanib (light blue), various doses of OM-153 (orange), or vehicle control (0.1% BSA, muted blue) (all 0.001% DMSO). Numbers in bold indicate fold activation (×) of positive controls relative to vehicle control. Boxplots show median, first and third quartiles, and whiskers (min–max) from five combined independent experiments (n = 3 replicates each). Stippled lines denote mean values of IPF-RC or TGFβ positive controls, set to 1. *P*-values, two-tailed t-test vs. respective positive controls.

Next, ECM synthesis of pro-collagen type III (PRO-C3), type VI (PRO-C6), and fibronectin (FBN-C) was evaluated using the Scar-in-a-Jar assay, a pseudo-3D NHLF model that captures essential features of fibrogenesis, including collagen deposition and ECM assembly, under physiologically relevant culture conditions for up to 12 days. Consistent with the 2D model, IPF-RC stimulation elevated PRO-C3, PRO-C6, and FBN-C protein levels relative to the control, whereas OM-153 concentration-dependently reduced all biomarkers without affecting cell viability (**figure 2a**, and **supplementary figure S2a**,**b**). By comparison, 100 nM nintedanib moderately suppressed these proteins, while a high dose of 1000 nM elicited more pronounced reductions (**figure 2b**).

**Figure 2.**
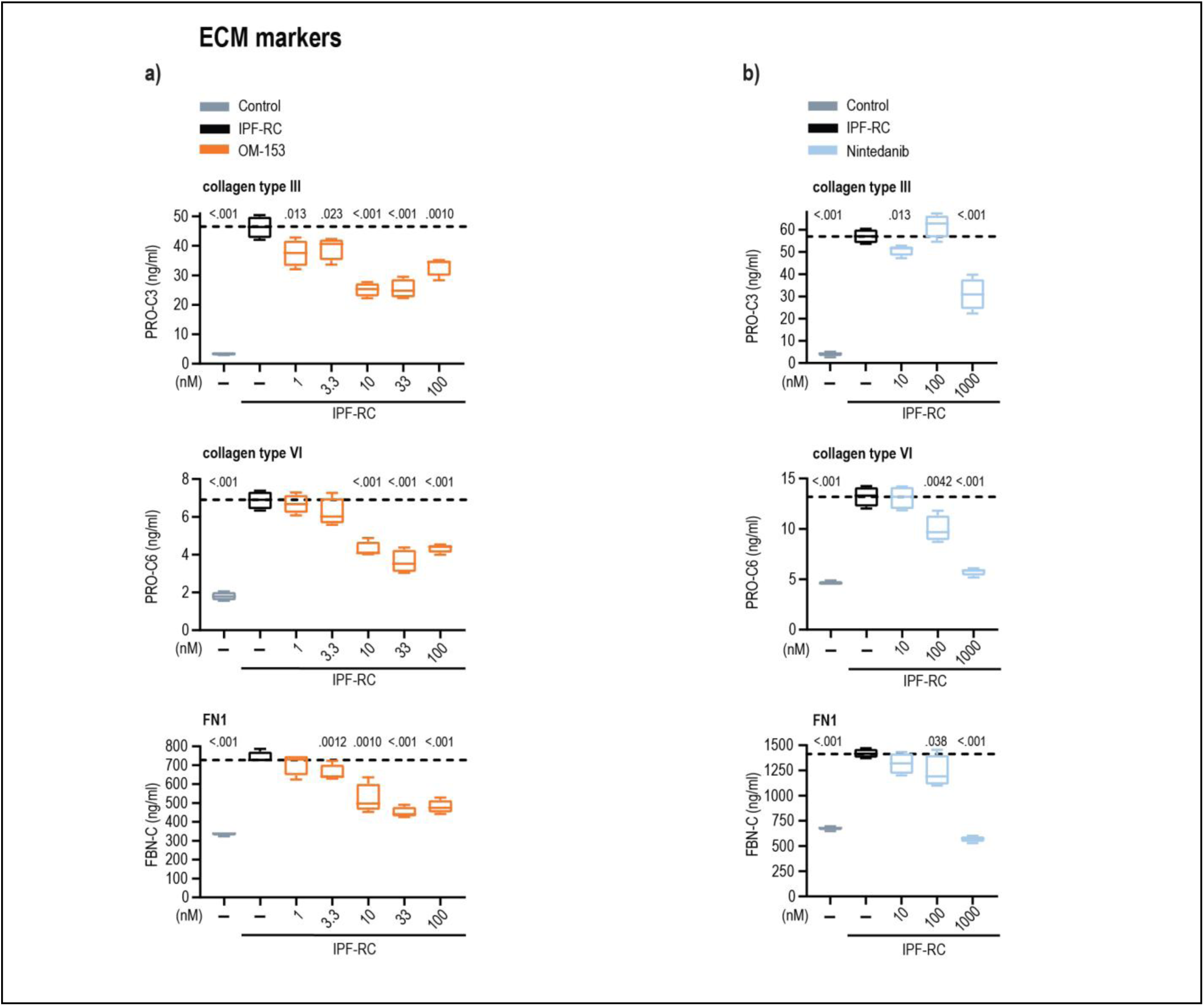
OM-153 reduces IPF-RC-stimulated ECM protein expression in normal human lung fibroblasts. **a**, **b)** 3D Scar-in-a-Jar assay of secreted pro-collagen type III (NordicPRO-C3™ [Pro-C3]), type VI (NordicPRO-C6™ [Pro-C6]), and fibronectin (NordicFBN-C™ [FBN-C]) (all in ng/mL) in NHLF supernatants after 12 days of stimulation with IPF-RC (1×, black) or vehicle control (0.1 ng/mL BSA, muted blue). Treatments: **a)** OM-153 (orange) or **b)** nintedanib (light blue) at the indicated doses under IPF-RC conditions (all 0.01% DMSO). Boxplots show median, first and third quartiles, and whiskers (min–max) from a representative dataset (three independent experiments, n = 4 replicates each). Stippled lines depict mean values of IPF-RC or TGFβ controls. *P*-values, one-tailed t-tests vs. IPF-RC control.

Collectively, these findings show that OM-153 potently and concentration-dependently suppresses ECM gene and protein expression in 2D and 3D lung fibroblast models under cytokine-driven, physiologically relevant IPF conditions.

### OM-153 decreases IPF-RC-stimulated ECM and FMT protein markers in normal human lung fibroblasts

FMT is a central pathogenic process in IPF, with myofibroblasts serving as the principal source of excessive ECM production [9]. To further assess the antifibrotic effects of OM-153, ECM deposition and the FMT marker actin alpha 2, smooth muscle (ACTA2) were examined in a 3D lung-on-a-chip model using NHLFs stimulated with IPF-RC or TGFβ. IPF-RC markedly increased collagen type I, fibronectin (FN1), and ACTA2 compared with control, whereas TGFβ induced these markers to a lesser extent (**figure 3a**). Both nintedanib and OM-153 reduced all three markers in IPF-RC–stimulated cells, while only collagen type I was significantly decreased under TGFβ stimulation (**figure 3a**). Consistently, immunoblot analysis in the 2D model revealed a concentration-dependent reduction of ACTA2 following OM-153 treatment in both IPF-RC– and TGFβ-stimulated NHLFs (**figure 3b**).

**Figure 3.**
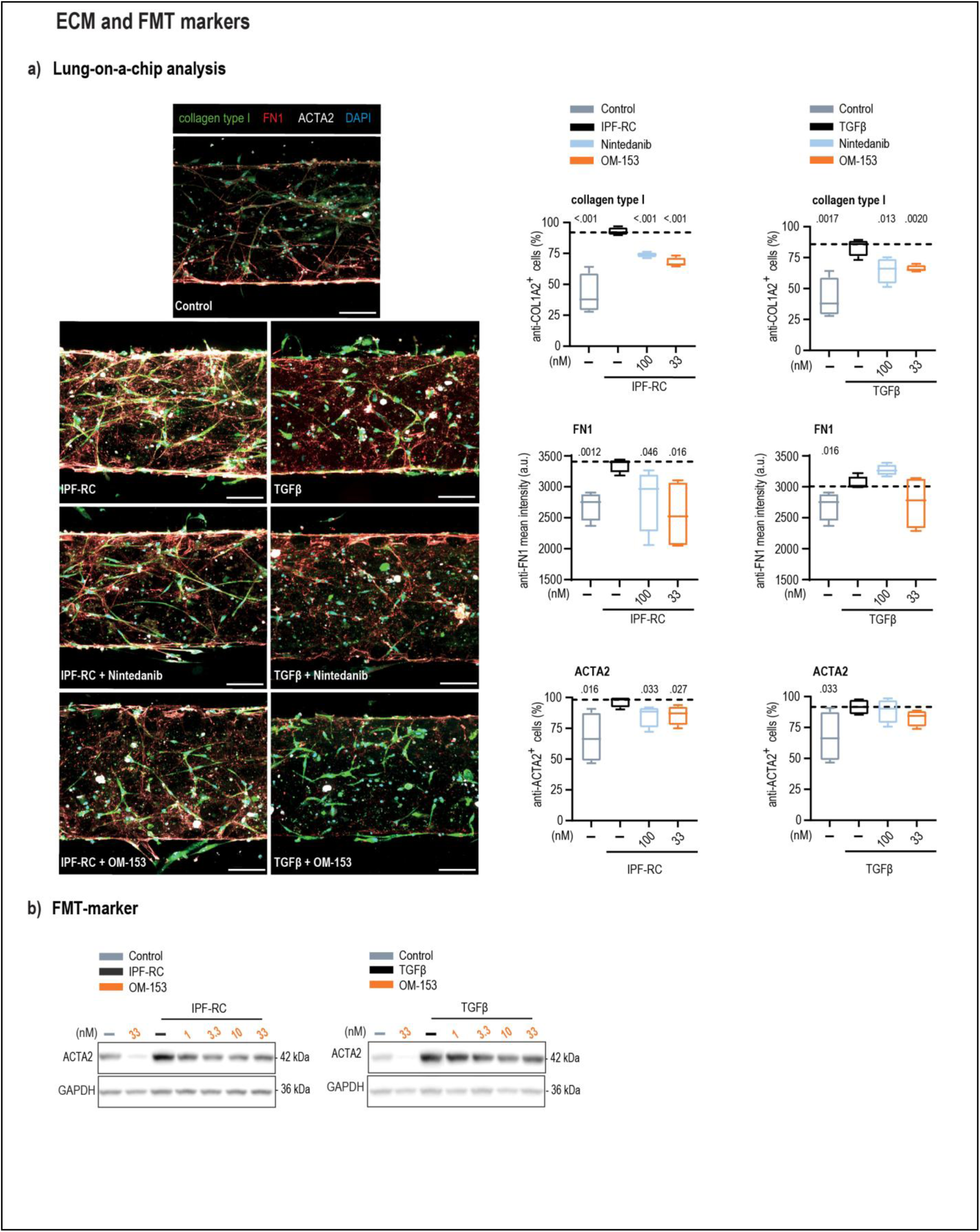
OM-153 decreases IPF-RC-stimulated ECM and FMT protein markers in normal human lung fibroblasts. **a)** Evaluation of ECM and FMT markers in 3D lung-on-a-chip model. Left, representative immunofluorescence images of collagen type I (green), FN1 (red), ACTA2 (white), and nuclei (DAPI, blue) in NHLFs. Cells stimulated for eight days with IPF-RC (1×) or TGFβ (0.3 ng/mL) (both, black) and treated with nintedanib (100 nM), OM-153 (33 nM), or control (0.1% BSA)(all in 0.001% DMSO). Scale bar, 100 µm. Right, quantification shown as the ratio of collagen type I- and ACTA2-positive cells to total nuclei, and FN1 mean fluorescence intensity for control (muted blue), IPF-RC (black), TGFβ (black), nintedanib (light blue), and OM-153 (orange). Boxplots show median, first and third quartiles, and whiskers (min–max) for combined data from four independent experiments. Stippled lines depict mean values of IPF-RC or TGFβ positive controls. *P*-values, one-tailed t-test vs. respective positive controls. **b)** Representative immunoblots of total protein extracts showing ACTA2 in NHLFs cultured in 2D, stimulated with IPF-RC (1×, left), TGFβ (0.3 ng/mL, right), or control (0.1% BSA)(all in 0.0003% DMSO), and treated with the indicated doses of OM-153 for 72 hours.

Together, these findings demonstrate that OM-153 effectively attenuates fibrogenesis and FMT in IPF-RC–stimulated NHLFs across *in vitro* models.

### OM-153 suppresses IPF-RC–induced profibrotic transcriptional programs in normal human lung fibroblasts

To characterize the broader transcriptional impact of IPF-RC stimulation and its modulation by OM-153, RNA sequencing was performed on NHLFs exposed to control medium, OM-153 (10 nM), IPF-RC, IPF-RC + OM-153 (10 nM), or IPF-RC + nintedanib (100 nM). A principal component analysis (PCA) demonstrated that OM-153 shifted the transcriptomic profile away from both control and IPF-RC clusters, whereas IPF-RC + nintedanib clustered closely with IPF-RC consistent with minimal transcriptomic modulation (**figure 4a**). DESeq2-generated volcano plots identified numerous differentially expressed genes (DEGs) for OM-153 vs control, IPF-RC vs control, and IPF-RC + OM-153 vs IPF-RC, while few significant DEGs were observed for IPF-RC + nintedanib vs IPF-RC, further underscoring its limited impact (**figure 4b**, **supplementary figure S4** and **table 1**).

**Figure 4.**
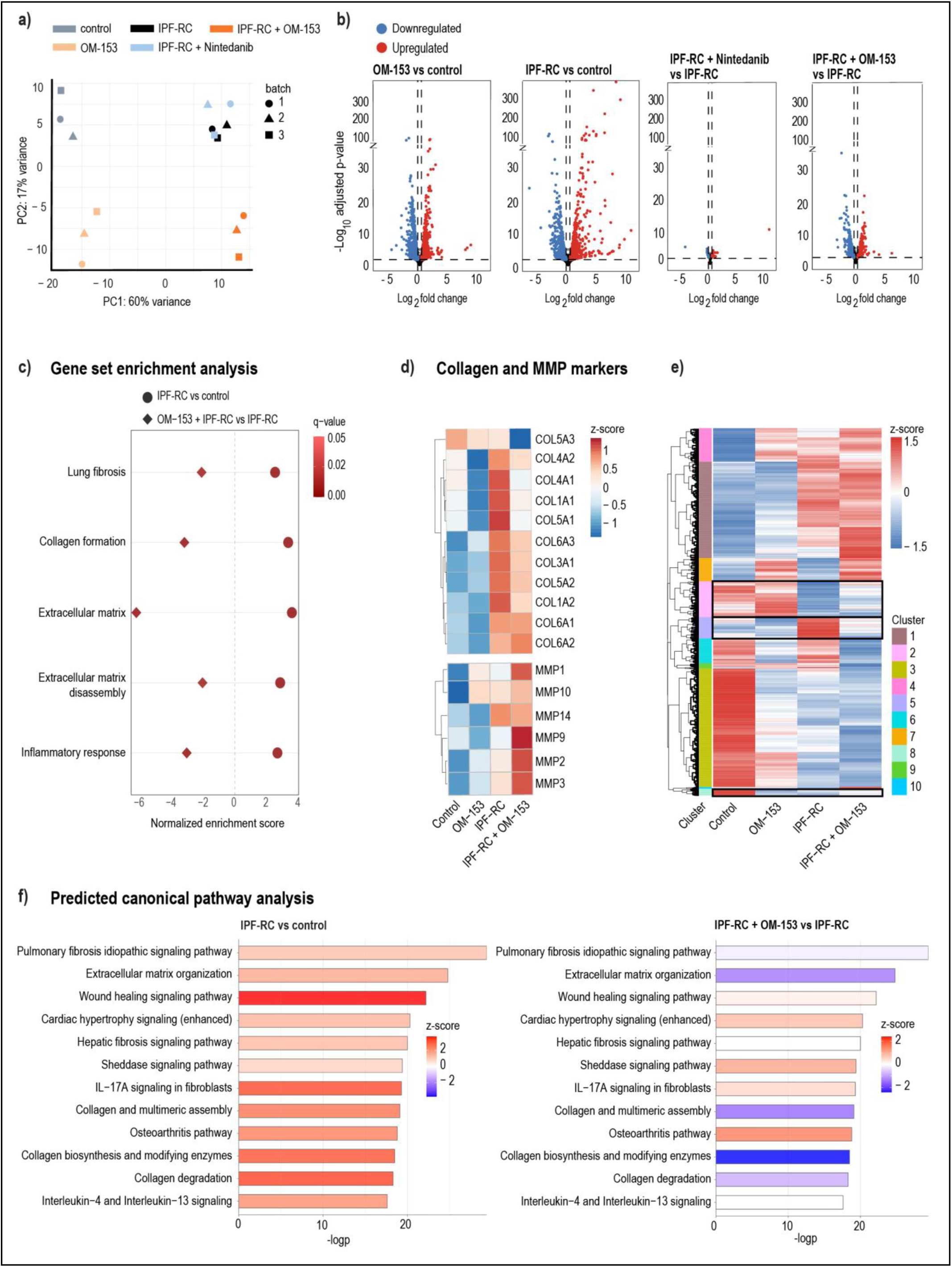
OM-153 suppresses IPF-RC-induced profibrotic transcriptional programs in normal human lung fibroblasts. **a)** PCA score plot showing gene expression diversity between groups. **a**-**e**) mRNA sequencing of NHLFs cultured for 72 hours (n = 3 per group). Experimental groups: Control (0.1% BSA), OM-153 (10 nM), IPF-RC (1×), IPF-RC + nintedanib (100 nM), and IPF-RC + OM-153 (10 nM)(all in 0.001% DMSO). **b)** Volcano plots of DEGs for the indicated comparisons: OM-153 vs. control, IPF-RC vs. control, IPF-RC + nintedanib vs. IPF-RC, and IPF-RC + OM-153 vs. IPF-RC. –log₁₀(p-value) plotted against log₂ fold change. Significantly downregulated (blue) and upregulated (red) genes shown (±0.3 fold change, *P* < 0.01). **c)** GSEA analysis of selected gene sets. Comparisons: IPF-RC vs. control (circles) and IPF-RC + OM-153 vs. IPF-RC (diamonds). Significant gene sets (q < 0.05) shown in a pink–red gradient, normalized enrichment scores indicate magnitude and direction (positive = upregulated, negative = downregulated). **d)** Hierarchical heatmap of transcriptional profiles (z-scores of log₂ fold change) for selected ECM and ECM-remodeling markers (collagens, MMPs) across control, OM-153, IPF-RC, and IPF-RC + OM-153. **e)** Hierarchical clustering heatmap of z-score expression for significant genes in IPF-RC vs. control and IPF-RC + OM-153 vs. IPF-RC. Ten gene clusters displayed: Clusters 2, 5, and 8 show a rescue effect by OM-153 in the IPF-RC condition (black boxes, detailed in **supplementary figure. S4**). Red indicates expression above the mean, blue below (z-score = 1.5 to –1.5). **f)** Canonical pathway enrichment predicted by IPA. Bar length indicates –log₁₀(*P-*value) for enrichment, color shows activation z-score based on DEGs in IPF-RC vs. control and IPF-RC + OM-153 vs. IPF-RC. Red (z-score > 0) indicates predicted activation, blue (z-score < 0) predicted inhibition. Pathways with *P*-values ≥ 0.05 or NaN z-scores were excluded.

Gene set enrichment analysis (GSEA) revealed significant positive enrichment of fibrosis-associated gene sets upon IPF-RC stimulation, which was reversed by OM-153, indicating suppression of fibrogenic transcriptional programs (**figure 4c**). IPF-RC upregulated collagens and ECM-remodeling matrix metalloproteinases (MMPs), while OM-153 broadly downregulated collagens and further upregulated MMPs (**figure 4d**), indicating an antifibrotic shift in ECM regulation.

Hierarchical clustering for the heatmap of DEGs from IPF-RC vs control and IPF-RC + OM-153 vs IPF-RC (**figure 4e**), followed by ingenuity pathway analysis (IPA), revealed activation of inflammation, IPF signaling, ECM organization, and collagen biosynthesis pathways by IPF-RC, all of which were reversed by OM-153 (**figure 4f** and **supplementary table 1**). Heatmap clusters 2, 5, and 8 specifically exhibited OM-153-mediated rescue of the IPF-RC transcriptional signature, with pathway analysis confirming this response profile (**supplementary figure 4a** and **table 1**).

Collectively, these data demonstrate that IPF-RC elicits a disease-relevant profibrotic program in NHLFs, and that OM-153 potently counteracts these transcriptional and pathway-level responses, supporting its potential as a targeted antifibrotic therapy.

### OM-153 inhibits IPF-RC-stimulated WNT/β-catenin and downregulates YAP target genes in normal human lung fibroblasts

OM-153 has been previously shown to inhibit WNT/β-catenin and YAP signaling, both of implicated in IPF progression [6]. First, to assess target engagement, stabilization of the direct TNKS substrates AXIN1 in the WNT/β-catenin signaling pathway and AMOTL1 in the YAP signaling pathway was examined. Immunoblotting demonstrated concentration-dependent stabilization of both proteins, confirming on-target activity (**Figure 5a**,**d**). In the WNT/β-catenin pathway, OM-153 reduced the transcriptionally active form of β-catenin (**figure 5a**). Transcriptomic profiling revealed that IPF-RC stimulation significantly upregulated WNT/β-catenin target genes, including the universal marker *AXIN2*, all of which were suppressed by OM-153, as confirmed by real-time qRT-PCR (**figure 5b**,**c**). In the YAP signaling pathway, OM-153 reduced nuclear accumulation of the transcriptional regulator YAP (**figure 5d**). Although IPF-RC did not upregulate selected YAP target genes, OM-153 significantly downregulated their expression, including *AMOTL2* and YAP-associated upstream regulators, as confirmed by pathway analysis (**figure 5e**,**f** and **supplementary table 1**).

**Figure 5.**
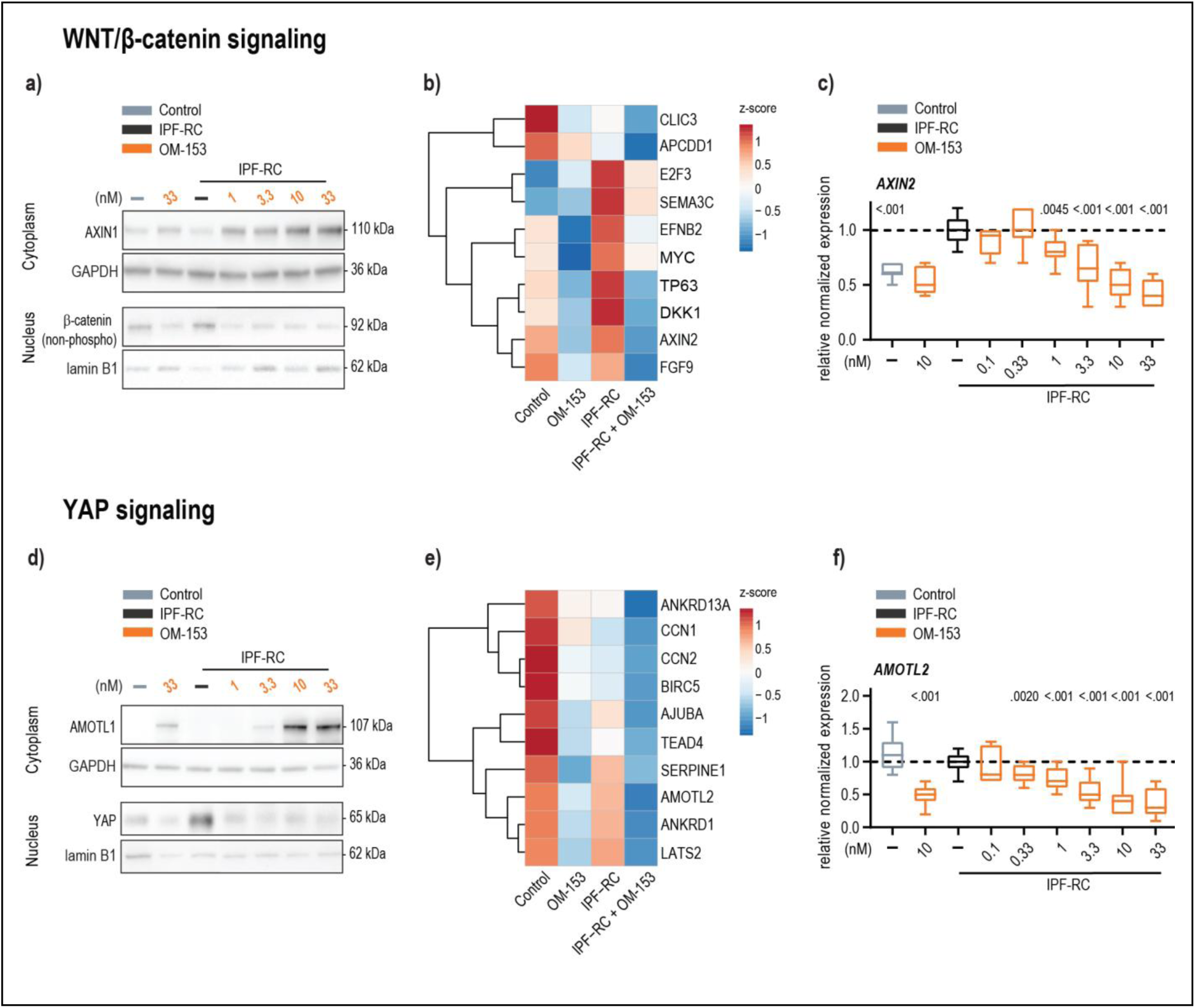
OM-153 inhibits IPF-RC-induced WNT/β-catenin signaling and downregulates YAP target genes in normal human lung fibroblasts. **a)** Representative immunoblots demonstrating TNKS target engagement via stabilization of cytoplasmic AXIN1, and reduction of the nuclear transcriptionally active form of β-catenin (non-phospho[Ser33/37/Thr41]). **a**-**f**) NHLFs were cultured for 72 hours under IPF-RC stimulation (1×) and treated with OM-153 at indicated doses or vehicle control (0.1% BSA)(all in 0.001% DMSO). **a**, **d**) GAPDH (cytoplasmic) and lamin B1 (nuclear) were used as loading controls. **b)** Hierarchical heatmap (z-scores of log₂ fold change) of selected WNT/β-catenin signaling target genes. **b**, **e)** Experimental groups: Control (0.1% BSA), OM-153 (10 nM), IPF-RC (1×), and IPF-RC + OM-153. **c)** Real-time qRT-PCR analysis of the WNT/β-catenin signaling target gene *AXIN2*. **c**, **f**) Boxplots show median, first and third quartiles, and whiskers (min–max) from five combined independent experiments (n = 3 replicates each). Stippled lines depict mean values of IPF-RC positive controls, set to 1. *P*-values, one-tailed t-test vs. IPF-RC positive controls. **d)** Representative immunoblots demonstrating TNKS target engagement via stabilization of cytoplasmic AMOTL1 and modulation of nuclear YAP and TAZ. **e)** Hierarchical heatmap (z-scores of log₂ fold change) of selected YAP-signaling target genes. **f)** Real-time qRT-PCR analysis of the YAP signaling target gene *AMOTL2*.

In conclusion, these findings demonstrate that IPF-RC selectively activates WNT/β-catenin but not YAP signaling in NHLFs, and that OM-153 potently inhibits WNT/β-catenin activation while suppressing YAP target gene expression.

### OM-153 reduces soluble collagen in BALF and collagen type I in a mouse bleomycin-induced lung fibrosis model

The bleomycin-induced lung fibrosis model is widely used in antifibrotic research, providing valuable insights into fibrotic mechanisms and biological drug effects [38]. To extend the *in vitro* findings, the antifibrotic efficacy of OM-153 was evaluated in this model. OM-153 was administered perorally at 10 mg/kg twice daily (PO-BID)[34], with nintedanib included as a comparator at 30 mg/kg once daily (PO-QD), starting on day 7 after bleomycin instillation.

At the experimental endpoint on day 21, histopathological analysis and Ashcroft scoring confirmed that bleomycin induced fibrosis and lung injury compared with controls (**figure 6a** and **supplementary figure S5b**). Ashcroft scoring of Masson’s trichrome staining (**figure 6a**), but not H&E (**supplementary figure S5b**), showed an effect approaching a statistically significance after treatment with OM-153 (*P* = 0.060) and nintedanib (*P* = 0.099). Bleomycin also increased infiltration of inflammatory immune cells, while tissue cytokine levels were not elevated (**supplementary figures S6a**,**b**). No changes in body weight were observed in any treatment group (**supplementary figure S5a**). Among treatments, only nintedanib reduced leukocyte and monocyte infiltration (**supplementary figure S6a**). Bleomycin challenge significantly increased soluble collagen in bronchoalveolar lavage fluid (BALF) and collagen type I in lung tissue, both of which were attenuated by OM-153 treatment, but not by nintedanib (**figure 6b** and **supplementary figure S7**). By contrast, the clinically relevant ECM biomarkers collagen type III, type VI, and FN1 were not induced (**figure 6b** and **supplementary figure S7**).

**Figure 6.**
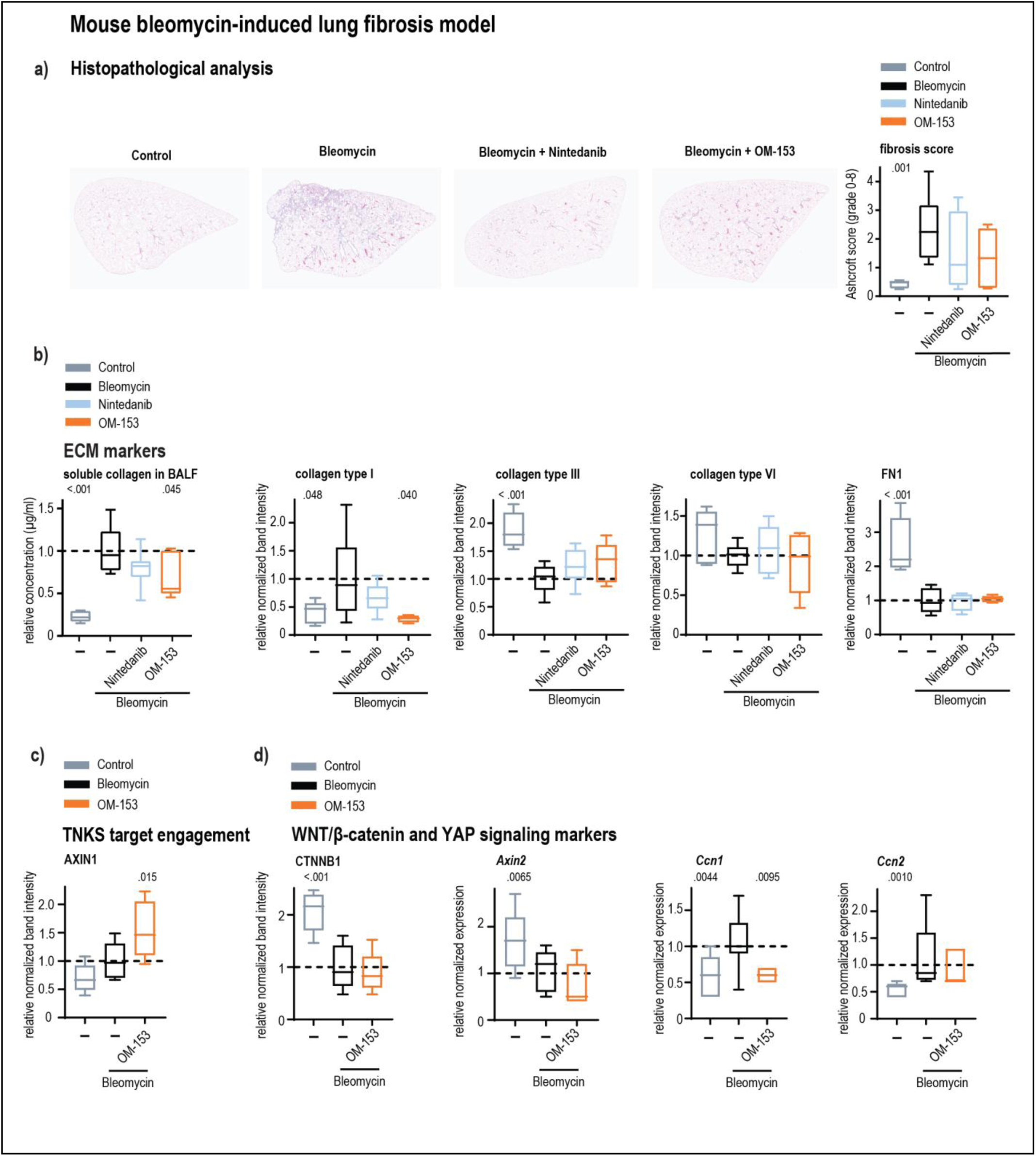
OM-153 reduces soluble collagen in BALF and collagen type I in a mouse bleomycin-induced lung fibrosis model. **a)** Histopathological analysis of lung tissue. Images of representative Masson’s Trichrome–stained sections (left) and quantification of fibrosis (Ashcroft score, grade 0–8, right). Scale bar: 100 µm. **a-d)**, C57BL/6 mice were challenged intratracheally (i.t.) with bleomycin (1.5 mg/kg) or control (n = 5, muted blue) on day 1. From day 7 through 20, mice received PO-BID treatment with vehicle (bleomycin, n = 8, black) or OM-153 (10 mg/kg, n = 5, orange), while nintedanib (30 mg/kg, n = 8, light blue) was administered PO-QD. Boxplots show median, first and third quartiles, and whiskers (min–max). Stippled lines denote mean bleomycin control values, set to 1. *P-*values, one-tailed t-test vs. bleomycin control. **b)** ECM protein analysis. Left, soluble collagen levels (µg/mL) in BALF. Right, relative quantification of immunoblots for lung collagen type I, type III, type VI, and FN1, normalized to GAPDH. **c)** TNKS target engagement. Relative quantification of AXIN1 immunoblots from lung tissue, normalized to GAPDH. **d)** Analysis of WNT/β-catenin and YAP signaling markers in lung tissue. Left, relative quantification of CTNNB1 immunoblots normalized to GAPDH. Right, real-time qRT-PCR of *Axin2*, *Ccn1*, and *Ccn2*.

To confirm target engagement, immunoblot analysis showed stabilization of AXIN1 following OM-153 treatment (**figure 6c** and **supplementary Figure S8**). Analysis of WNT/β-catenin signaling revealed no bleomycin-induced upregulation of CTNNB1 protein or *Axin2* mRNA. Conversely, the YAP target genes *Ccn1* and *Ccn2* were elevated, but with only *Ccn1* significantly reduced by OM-153 (**figure 6d** and **supplementary figure S8**).

Despite its limitations in activating key ECM markers and WNT/β-catenin signaling, the bleomycin-induced lung fibrosis model demonstrated that OM-153 stabilized AXIN1 and attenuated soluble collagen accumulation in BALF as well as type I collagen deposition in lung tissue, thereby confirming *in vivo* target engagement and antifibrotic activity.

### OM-153 suppresses ECM and FMT markers and inhibits WNT/β-catenin and YAP signaling in *ex vivo* human lung fibrosis models

To establish human proof-of-concept for TNKS inhibitor efficacy, PCLS were employed as an *ex vivo* model of IPF. This system preserves the full cellular complexity of the lung and provides a physiologically relevant platform for drug evaluation. PCLS were prepared from non-PF donor tissue and stimulated with IPF-RC to model fibrogenesis, as well as from end-stage PF donor tissue, followed by treatment.

In PCLS from non-PF donors, real-time qRT-PCR analysis demonstrated that IPF-RC stimulation induced a significant upregulation of *COL1A1*, the clinically relevant ECM markers *COL3A1*, *COL6A1*, and *FN1* (**figure 7a**), as well as the FMT marker *ACTA2* (**figure 7c**), relative to control. These results indicate that IPF-RC can recapitulate the pro-fibrotic environment of human IPF lungs and activate key markers of fibrogenesis. OM-153 treatment suppressed the IPF-RC-induced expression of all markers, whereas the SoC compounds pirfenidone and nintedanib reduced most markers (**figure 7a**,**c**), all without impacting tissue viability (**supplementary figure S9a**). In PCLS from PF donors, OM-153 similarly reduced expression of these four markers, while the SoC compounds remained largely ineffective (**figure 7b**,**d** and **supplementary figure S9b**).

**Figure 7.**
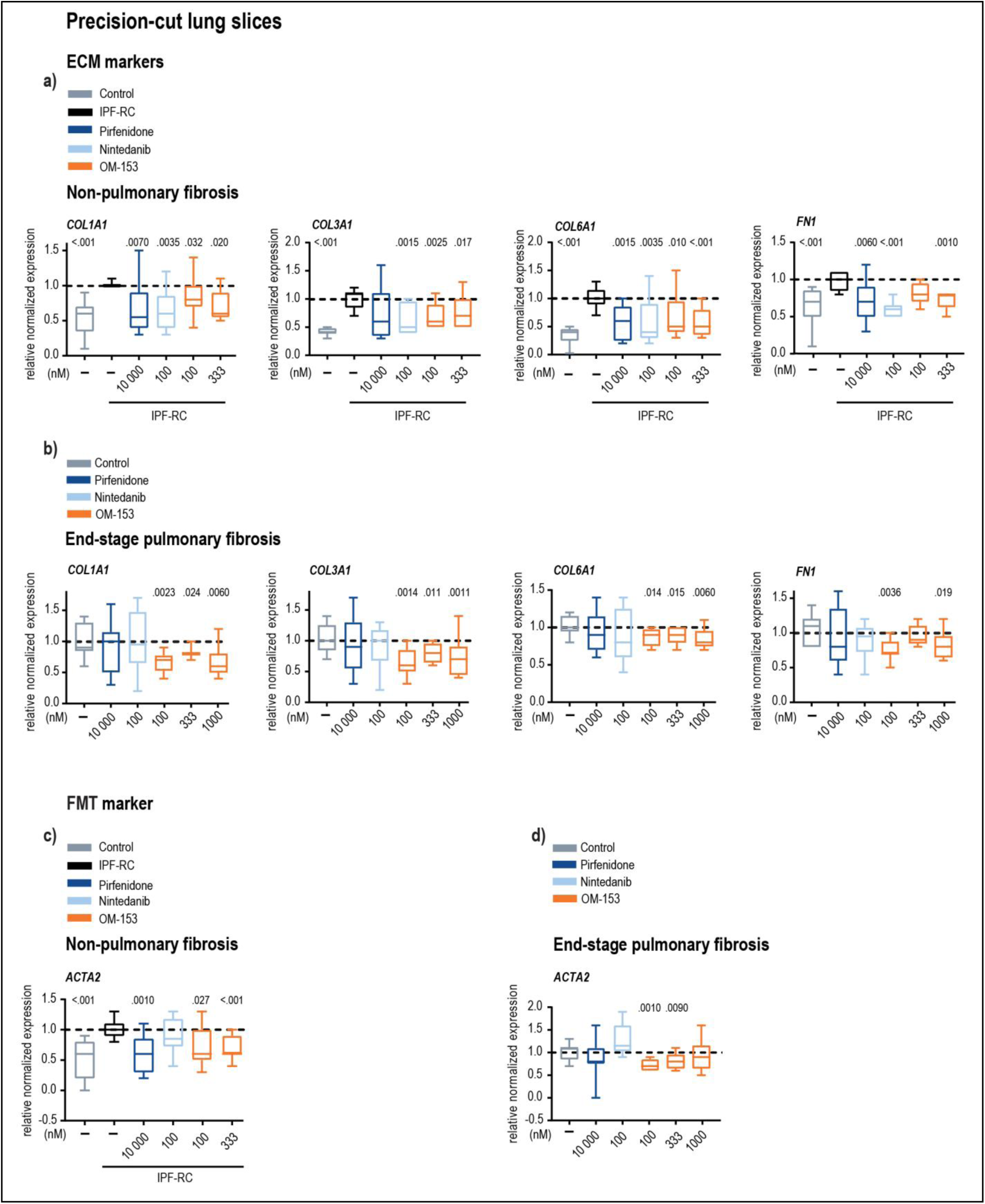
OM-153 suppresses ECM and FMT markers in *ex vivo* human lung fibrosis models. **a**, **b)** Real-time qRT-PCR analysis of ECM markers (*COL1A1*, *COL3A1*, *COL6A1*, and *FN1*) in PCLS from **a**) non-PF and **b**) PF donors. **a-d)** PCLS (n = 3 non-PF or PF donors, six slices per donor) were cultured for 72 hours. Non-PF PCLS were stimulated with IPF-RC (3×, 0.01% DMSO, black). All slices were treated with the indicated doses of pirfenidone (deep blue), nintedanib (light blue), OM-153 (orange), or control (medium, muted blue). Boxplots show median, first and third quartiles, and whiskers (min–max) for combined data from three donors (six PCLS each), pooled in pairs, with three replicates each. Stippled lines indicate mean IPF-RC control (for non-PF) or control values (for PF) set to 1. *P*-values, one-tailed t-test vs. respective controls. **c**, **d)** Real-time qRT-PCR analysis of the FMT marker *ACTA2* in PCLS from **c)** non-PF and **d)** PF donors.

To determine effects on WNT/β-catenin and YAP signaling pathways, immunoblotting and real-time qRT-PCR revealed that OM-153 stabilized the direct TNKS target AXIN1 and significantly suppressed expression of the WNT/β-catenin target gene *AXIN2* in both donor groups (**figures 8a**,**b** and **supplementary figure S10a**,**b**). OM-153 also reduced expression of the YAP target genes *AMOTL2*, *CCN1*, and *CCN2*, with the exception of *CCN2* in non-PF tissue (**figures 8c**,**d**). In addition, and consistent with *in vivo* findings, OM-153 did not reduce inflammatory cytokine levels in either non-PF or PF PCLS (**supplementary figures S11a**,**b**).

**Figure 8.**
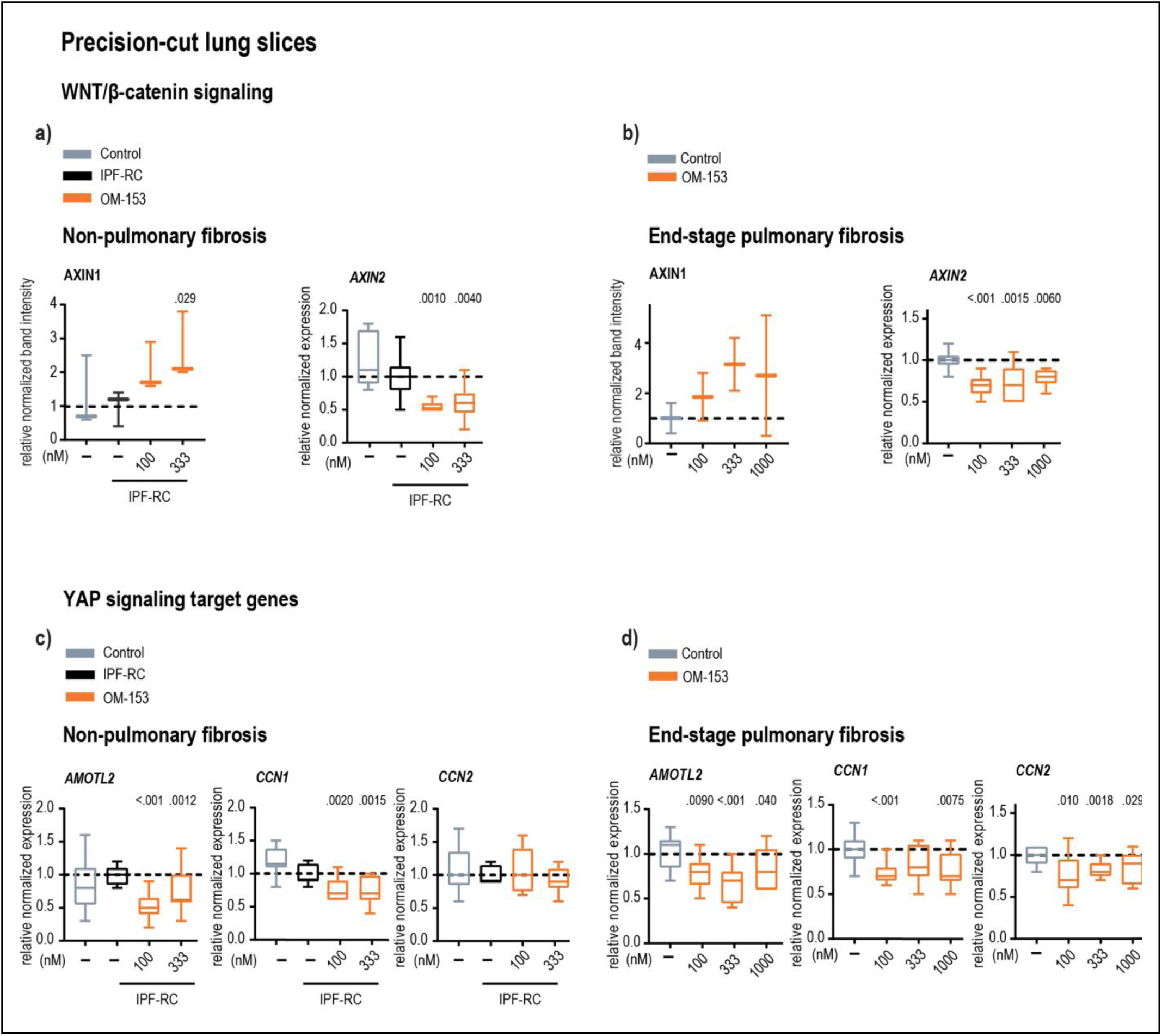
OM-153 inhibits WNT/β-catenin and YAP signaling in *ex vivo* human lung fibrosis models. **a**, **b)** Evaluation of WNT/β-catenin signaling markers in PCLS from **a**) non-PF and **b**) PF donors. Left, relative quantification of immunoblots showing the TNKS target engagement marker AXIN1, normalized to GAPDH. Right, real-time qRT-PCR analysis of the WNT/β-catenin signaling target gene *AXIN2*. **a-d)** PCLS (n = 3 non-PF or PF donors, six slices per donor) were cultured for 72 hours. Non-PF PCLS were stimulated with IPF-RC (3×, 0.01% DMSO, black). All slices were treated with the indicated doses of OM-153 (orange) or control (medium, muted blue). Boxplots show median, first and third quartiles, and whiskers (min–max) for combined data from three donors (six PCLS each), pooled in pairs, with three replicates each. Stippled lines indicate mean IPF-RC control (for non-PF) or control values (for PF), set to 1. *P*-values, one-tailed t-test vs. respective controls. **c**, **d)** Real-time qRT-PCR analysis of YAP signaling target genes *AMOTL2*, *CCN1*, and *CCN2* in **c**) non-PF and **d**) PF donors.

In conclusion, these findings establish human proof-of-concept for the efficacy of TNKS inhibition. OM-153 effectively suppresses fibrogenic and FMT markers while modulating WNT/β-catenin and YAP signaling in both IPF-RC–stimulated non-PF and PF PCLS, highlighting its potential as a targeted antifibrotic therapy in complex human lung tissue.

## Discussion

Here, we demonstrate that the selective TNKS inhibitor OM-153 stabilizes AXIN and AMOT, thereby promoting β-catenin degradation and cytoplasmic sequestration of YAP, resulting in dual suppression of WNT/β-catenin and YAP signaling. The treatment effectively counteracts FMT and pathological ECM synthesis and deposition across key preclinical models. *In vitro*, OM-153 suppressed cytokine-driven fibrogenesis in primary human lung fibroblasts under both 2D and 3D culture conditions. *In vivo*, OM-153 reduced collagen deposition in bleomycin-challenged mice, and *ex vivo* it attenuated ECM gene expression in PCLS derived from both cytokine-stimulated non-PF and end-stage PF donors.

IPF progression is driven by a complex cytokine network, yet most *in vitro* studies rely exclusively on TGFβ stimulation, limiting the evaluation of antifibrotic mechanisms that act independently of this pathway [4, 12, 39]. In NHLFs, IPF-RC robustly upregulated the clinically relevant collagens *COL3A1* and *COL6A1*, as well as *COL1A1*, whereas TGFβ alone induced only *COL1A1* and *COL3A1*. OM-153 markedly suppressed IPF-RC–induced expression of all three ECM genes, while neither OM-153 nor nintedanib altered TGFβ-stimulated cultures. In the lung-on-a-chip model, IPF-RC robustly increased ECM deposition and ACTA2 expression, whereas TGFβ caused comparatively moderate responses. Consistently, IPF-RC triggered a transcriptional program characteristic of active fibrogenesis, including activation of WNT/β-catenin signaling in NHLFs, and in addition upregulation of ECM markers in non-PF PCLS. Together, these findings underscore the value of IPF-RC as a physiologically relevant stimulus for evaluating antifibrotic biotargets such as TNKS acting beyond the TGFβ axis.

Importantly, TNKS inhibition exerts pleiotropic effects by modulating multiple signaling cascades in parallel, including WNT/β-catenin and YAP signaling [13, 14]. Such coordinated pathway modulation may confer therapeutic benefit in IPF, where WNT/β-catenin, YAP, and TGFβ pathways are highly interconnected [6]. The multimodal activity of established and emerging antifibrotic agents, including nintedanib, pirfenidone, nerandomilast, and treprostinil, has been proposed to underlie their clinical efficacy despite limited pathway selectivity [2, 40]. OM-153 represents a mechanistically distinct yet comparably pleiotropic approach, positioning TNKS inhibition as a promising strategy for broad suppression of fibrogenic signaling in IPF, either as monotherapy or in combination therapies. Future antifibrotic therapies will likely be combined with orally administered SoC agents associated with gastrointestinal adverse effects [4]. Hence, alternative routes of administration, such as inhaled delivery, may reduce systemic exposure, minimize drug–drug interactions, and enhance pulmonary efficacy [4].

OM-153 did not significantly affect fibroblast proliferation *in vitro*, or compromise tissue viability as indicated by LDH-release assays in PCLS. Similarly, no adverse effects were observed in the lungs following oral administration in the bleomycin model or in a 28-day repeated dose toxicity study [34]. Clinical data using the TNKS inhibitor STP1002 have demonstrated systemic tolerability [35]. Nevertheless, a comprehensive evaluation of the therapeutic window and additional cell type–specific effects, including those on epithelial cell differentiation, dysfunctional pro-fibrotic transitional epithelial states, and immune populations such as macrophages, remains warranted [7–9].

In conclusion, our findings identify TNKS inhibition as a mechanistically distinct antifibrotic strategy that achieves coordinated suppression of WNT/β-catenin and YAP signaling. The efficacy of OM-153 across complementary preclinical models supports further translational development, including inhalation-based delivery and combination strategies, to advance TNKS inhibition toward clinical application in IPF.

## Acknowledgements

We thank Stine Johansen and Maja Strangeways for technical assistance (both Nordic Bioscience). The authors used ChatGPT Business (OpenAI, GPT-5; accessed [13 October 2025– 04 November 2025]) for minor language and code editing. All outputs were reviewed and approved by the authors, who take full responsibility for the content.

## Author contributions

**S.A.B.** contributed to conceptualization, investigation, formal analysis, methodology, data interpretation, experimental work, supervised the project, and wrote the original manuscript draft. **I.J.** conducted 2D *in vitro* experiments and performed protein/mRNA analysis *in vivo* and *ex vivo* analysis, and contributed to methodology and validation. **M.F.S.**, **O.V.B.** performed bioinformatic analysis of RNA sequencing data, and contributed to visualization, methodology, data interpretation, and manuscript writing. **K.M.F.**, **M.C.L.**, **M.T.M.**, **I.L.M.** contributed to *in vitro* experiments, protein/mRNA analysis, validation, and methodology. **L.W.**, **C.H.** performed human PCLS data acquisition and analysis, as well as contributing to validation, visualization, and methodology. **D.J.**, **P.Z.**, **H.G.F.** provided access to human primary lung tissue and performed pathological diagnosis. **M.A.S.**, **M.A.K.** performed scar-in-a-jar assays, and contributed to data analysis and methodology. **T.L.**, **R.V.**, **E.F.**, **P.O.** carried lung-on-a-chip assays and contributed to data analysis and methodology. **S.J.**, **J.L.**, **H.H.** carried out experiments evaluating inflammation in animals, and contributed to data analysis, data interpretation and methodology. **S.R.** processed RNA raw data for bioinformatic analysis. **A.W., S.C.** contributed to conceptualization, investigation, methodology and data interpretation. **S.K.** contributed to conceptualization, investigation, methodology, data interpretation, and provided funding. **J.W.** contributed to conceptualization, investigation, formal analysis, methodology, data interpretation, experimental work, supervised the project, provided funding, wrote the original manuscript draft, and managed the project. **All** authors read and approved the final manuscript.

