## Supplementary figures for "Tankyrase inhibition demonstrates anti-fibrotic effects in preclinical pulmonary fibrosis models"

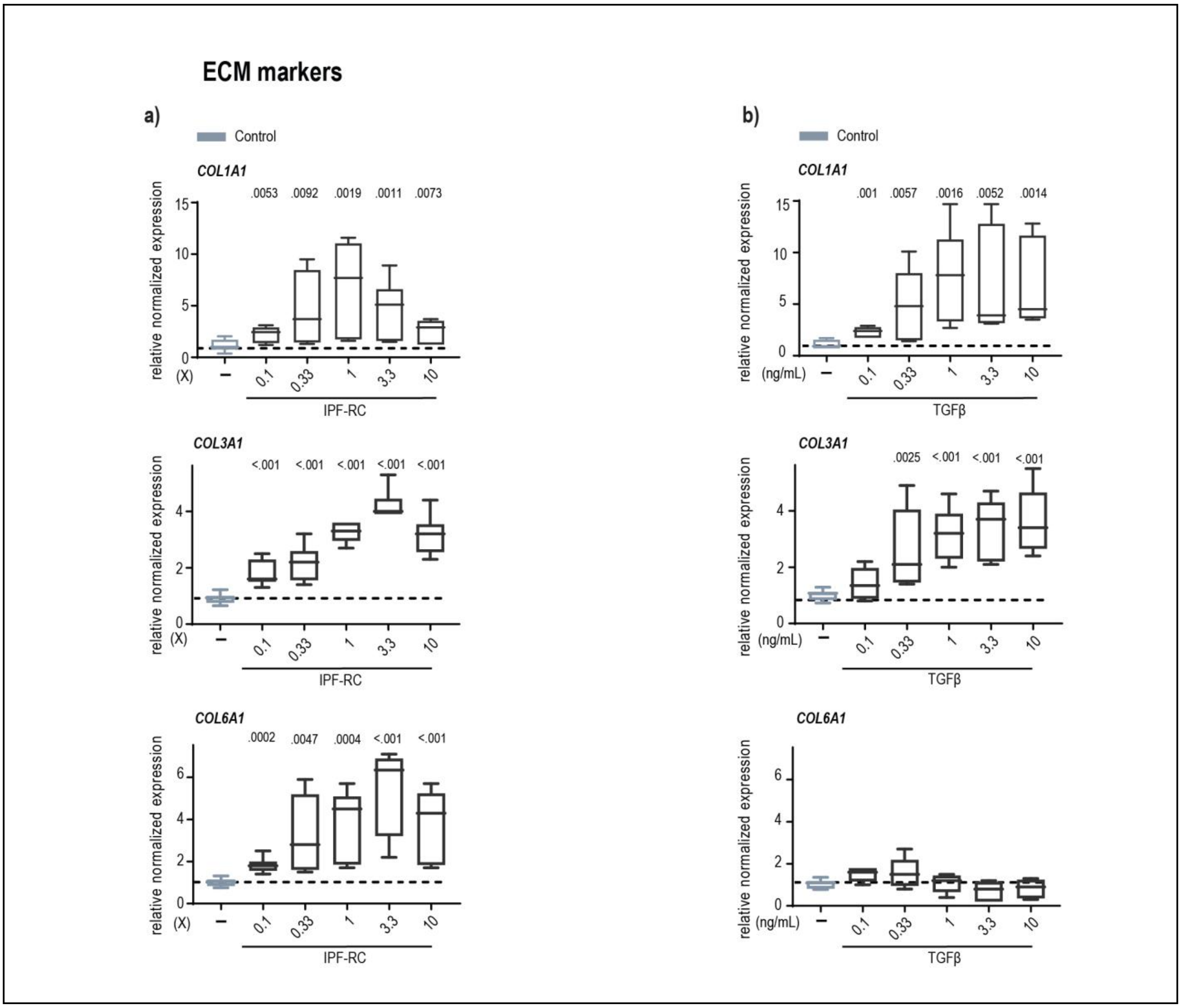


**Supplementary Figure S1. Comparison of ECM induction by IPF-RC versus TGFβ**

Real-time qRT-PCR analysis of ECM markers (*COL1A1*, *COL3A1*, *COL6A1*) in NHLFs cultured in 2D for 72 hours. Cells were stimulated with various concentrations of **a)** IPF-RC (0.1–10×) or **b)** TGFβ (0.1–10 ng/mL) and vehicle control (0.1% BSA, muted blue). Boxplots show median, first and third quartiles, and whiskers (min–max) for combined data from three independent experiments (n = 3 replicates each). Stippled lines denote mean vehicle control values, set to 1. *P*-values, two-tailed t-test vs. control.

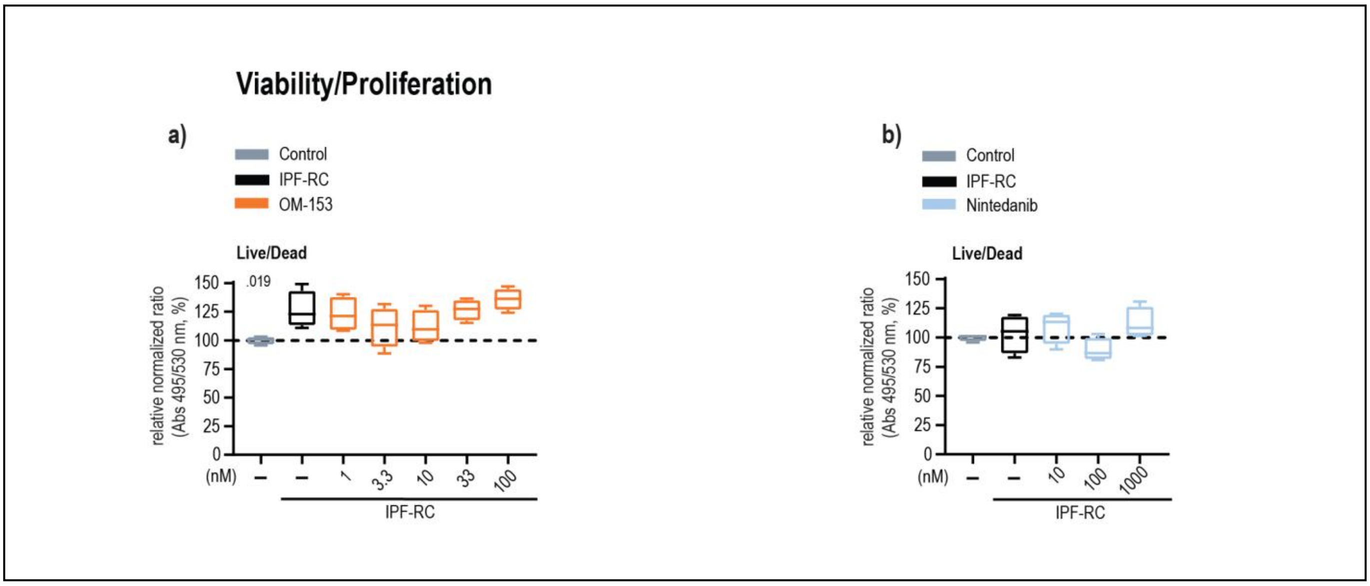


**Supplementary Figure S2. OM-153 viability assay in Scar-in-a-Jar in normal human lung fibroblasts.**

3D Scar-in-a-Jar endpoint analysis of cell viability (live/dead) from the experiment described in **figure 2**. Boxplots show median, first and third quartiles, and whiskers (min–max) from a representative dataset (three independent experiments, n = 4 replicates each). Stippled lines depict mean vehicle control values normalized to 100%. *P*-values, two-tailed t-test vs. IPF-RC control.

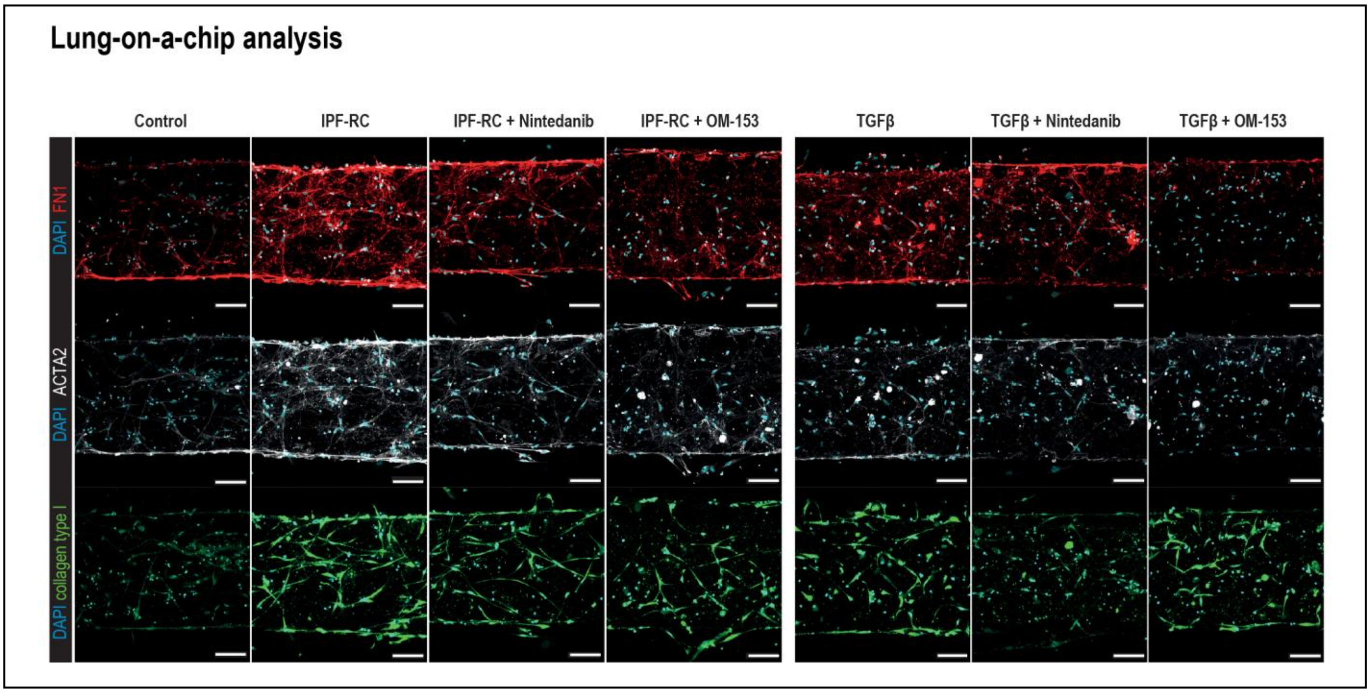


**Supplementary Figure S3. OM-153 decreases IPF-RC-stimulated ECM and FMT protein markers in normal human lung fibroblasts**

Evaluation of ECM and FMT markers in 3D lung-on-a-chip model, and representative individual immunofluorescence images corresponding to the merged images described in **figure 3a**. Collagen type I (green), FN1 (red), ACTA2 (white), and nuclei (DAPI, blue). Scale bar, 100 µm.

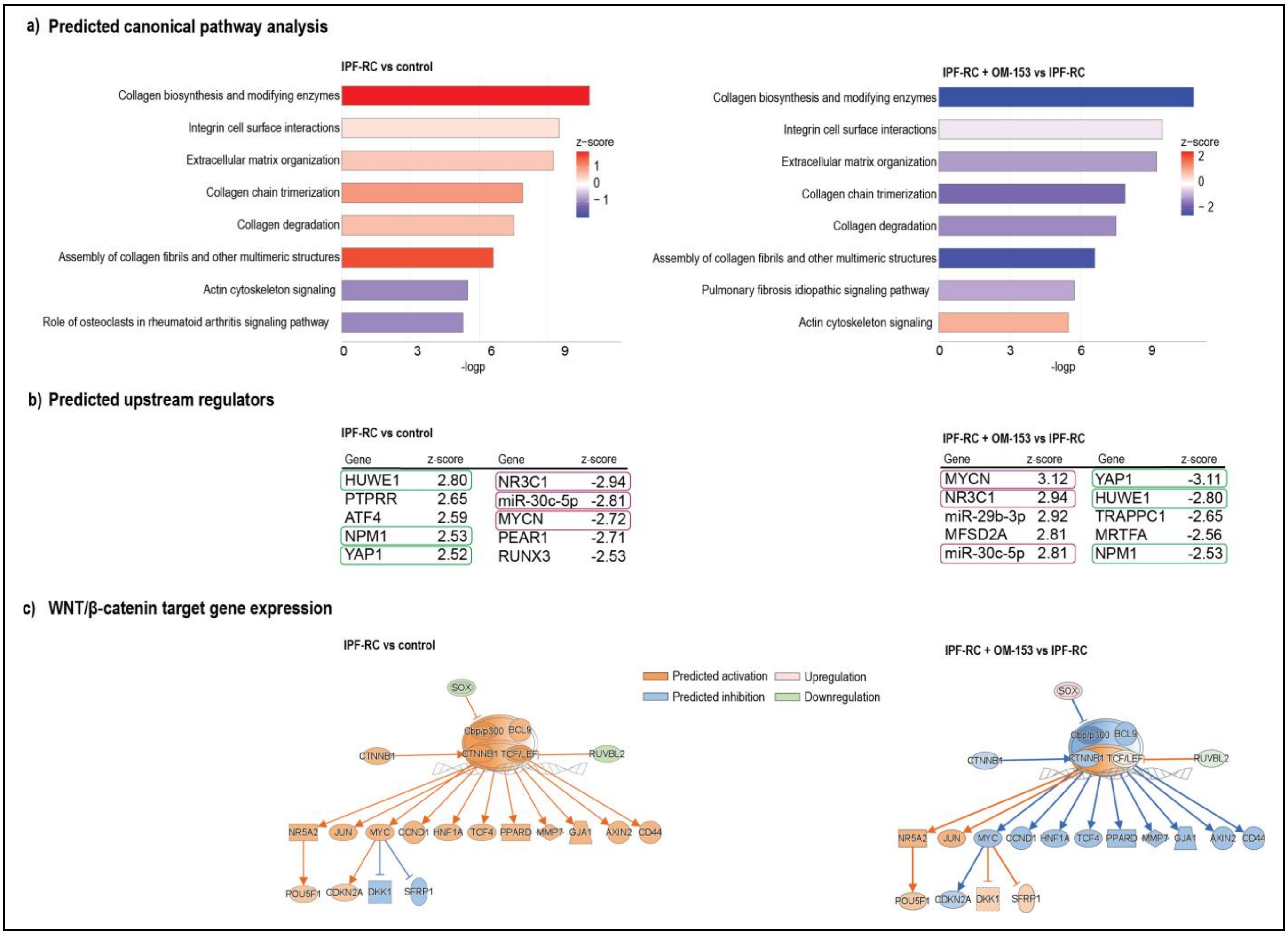


**Supplementary Figure S4 OM-153 suppresses IPF-RC-induced profibrotic transcriptional programs in normal human lung fibroblasts**

IPA performed using the combined gene list of clusters 2, 5, and 8 described in **figure 4e** for IPF-RC vs. control (left) and IPF-RC + OM-153 vs. IPF-RC (right).

**a)** Predicted pathway enrichment displayed (bar length = $-\log_{10}(P)$; bar color = activation z-score). Pathways with $P$-values $\geq 0.05$ were excluded.

**b)** Predicted upstream regulators showing the top and bottom five z-scored genes (purple and green boxes indicate reversals between contrasts). Molecule type: genes, RNAs, and proteins. Upstream regulators with $P$-values $\geq 0.05$ or absolute z-scores $< 2$ were excluded.

**c)** WNT/β-catenin pathway target gene expression (orange = predicted activation, red = upregulation, blue = predicted inhibition, green = downregulation; ±0.3 fold change, $P < 0.01$).

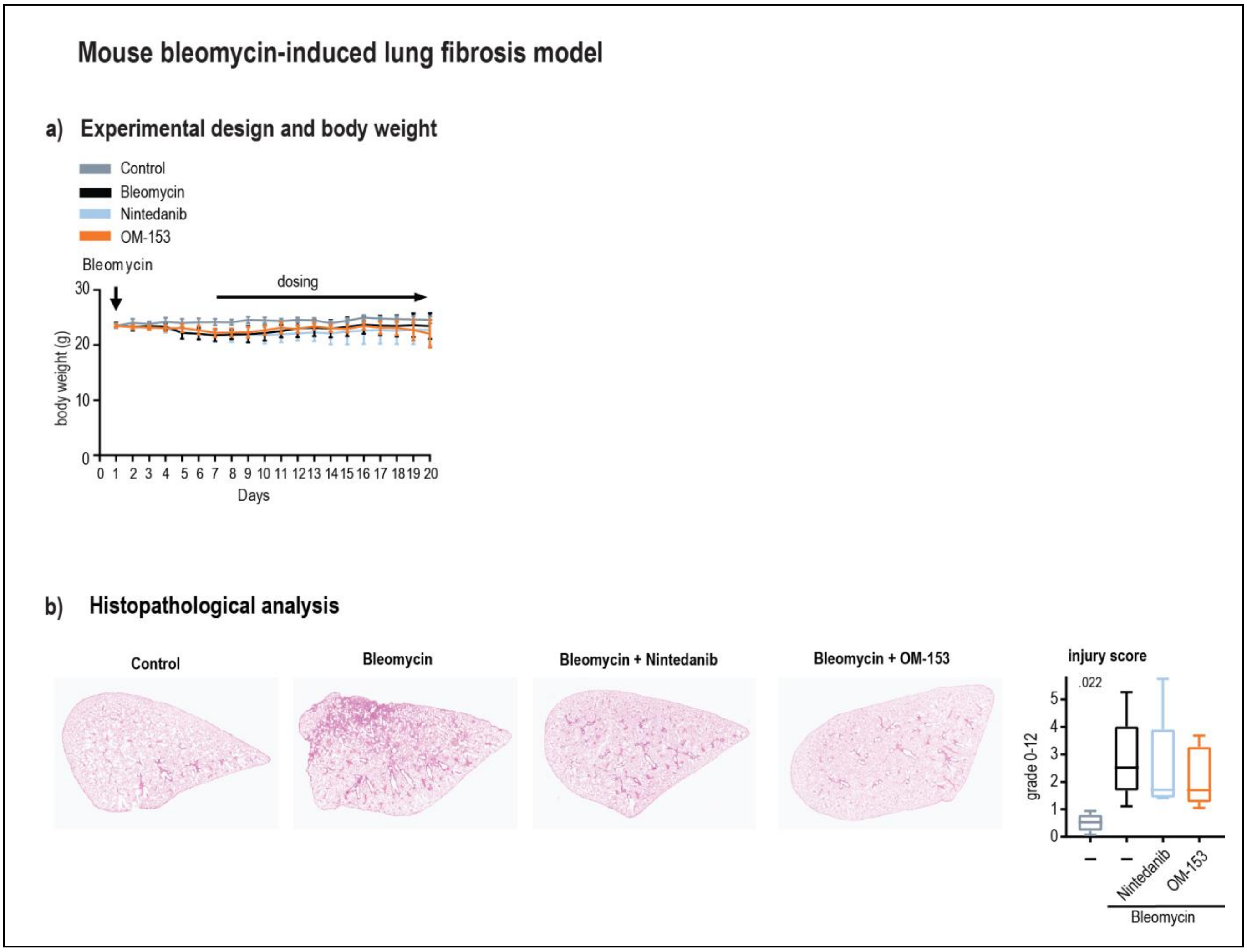


**Supplementary Figure S5. Experimental design, body weight, and Ashcroft score in a mouse bleomycin-induced lung fibrosis model**

**a**, **b)** Data from the experiment described in **figure 6**. **a)** Experimental design showing treatment regimens and body weight (g) from day 1 to day 20. Standard deviations are shown. No significant changes were documented using two-tailed t-tests vs. vehicle control.

**b)** Histopathological analysis of lung tissue showing hematoxylin–eosin (H&E)–stained section (left, scale bar: 100 µm.) and quantification of injury scores (Ashcroft score, grade 0–12; right). Boxplots show median, first and third quartiles, and whiskers (min–max).

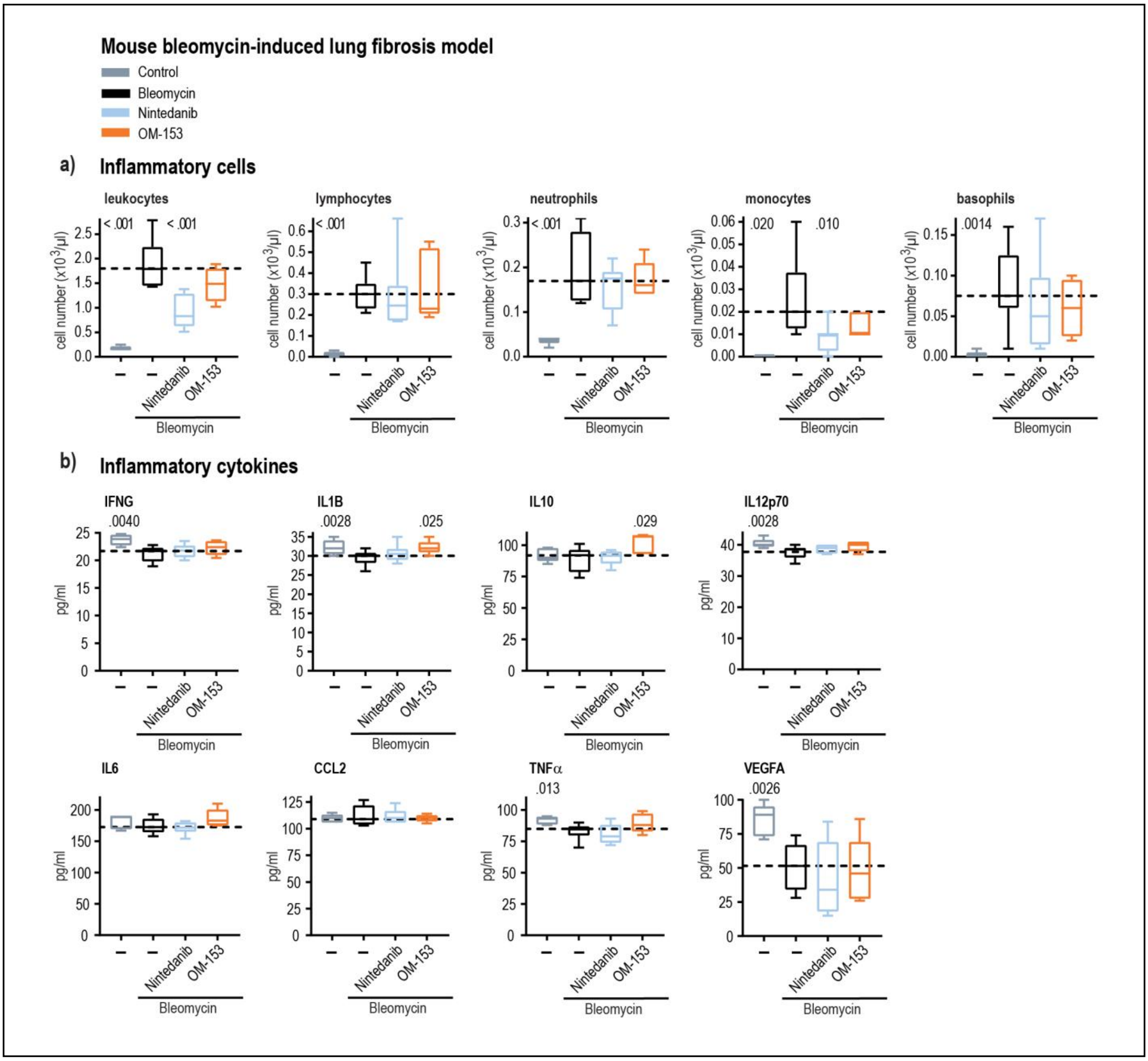


**Supplementary Figure S6. Inflammatory cells and cytokines in a mouse bleomycin-induced lung fibrosis model**

**a**, **b)** Data from the experiment described in **figure 6.** Boxplots show median, first and third quartiles, and whiskers (min–max). Stippled lines denote mean bleomycin control values. *P*-values, two-tailed t-test vs. bleomycin control. **a)** Inflammatory cells (leukocytes, lymphocytes, neutrophils, monocytes, and basophils) shown as cell number (×10³/µL). **b)** Inflammatory cytokines in lung extracts with mean concentrations (pg/mL).

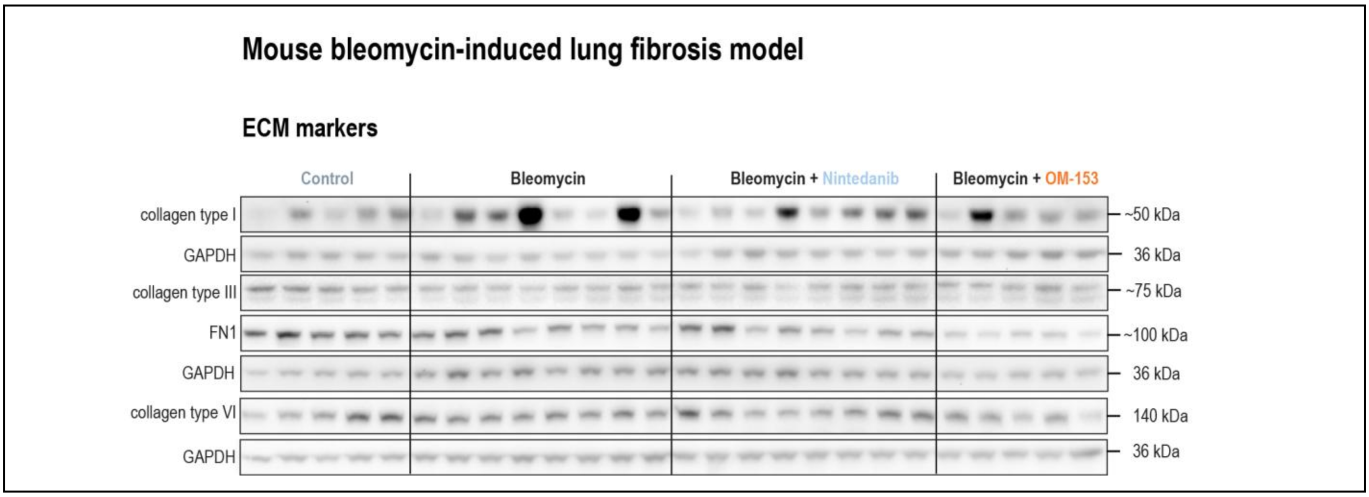


**Supplementary Figure S7. OM-153 reduces collagen type I in a mouse bleomycin-induced lung fibrosis model**

Representative immunoblots of ECM markers (collagen type I, collagen type III, FN1, and collagen type VI) in total lung extracts obtained from the experiment described in **figure 6b**. GAPDH was used as a loading control.

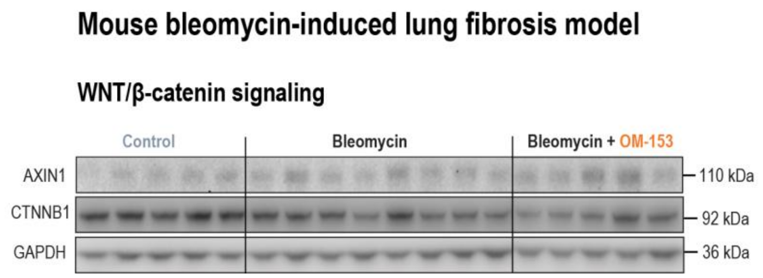


**Supplementary Figure S8. Immunoblot analysis of AXIN1 and CTNNB1 in a mouse bleomycin-induced lung fibrosis model**

Representative immunoblots of the TNKS target engagement marker AXIN1 and CTNNB1 in the WNT/β-catenin signaling pathway. The analysis was performed on total lung extracts obtained from the experiment described in **figure 6c**,**d**. GAPDH was used as a loading control.

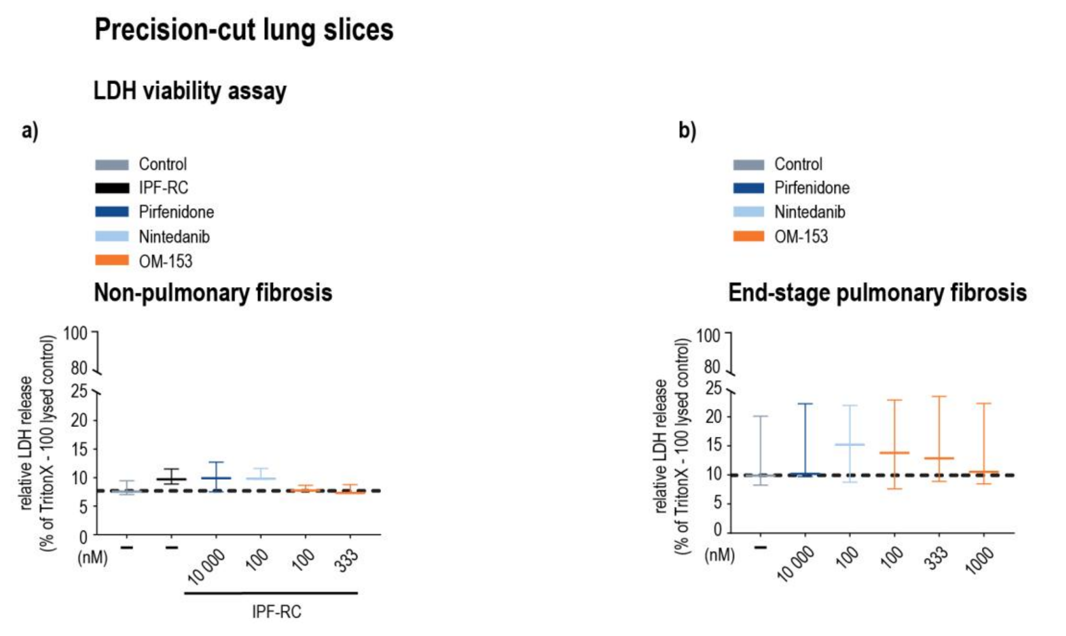


**Supplementary Figure S9. OM-153 does not affect tissue viability in *ex vivo* human lung fibrosis models**

**a**, **b)** Lactate dehydrogenase (LDH) release measured as a marker of tissue viability and presented as relative LDH release (% of Triton X-100–lysed control). Culture supernatants collected after 72 hours from the experiment described in **figure 7**. Boxplots show median, first and third quartiles, and whiskers (min–max) for combined data from three donors (six PCLS each), pooled in pairs. Stippled lines indicate mean vehicle control values. No significant changes were documented using two-tailed t-tests vs. control. **a)** PCLS from non-PF donor tissue, and **b)** PCLS from PF donor tissue.

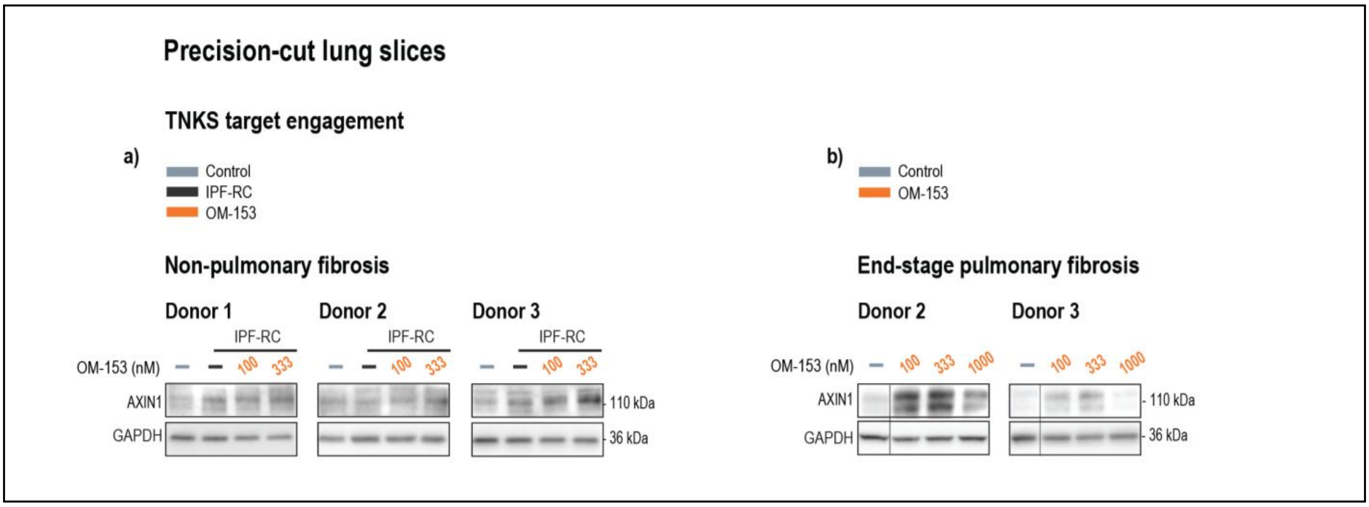


**Supplementary Figure S10. Immunoblot analysis of AXIN1 in *ex vivo* human lung fibrosis models**

**a**, **b)** Representative immunoblots of the TNKS target engagement marker AXIN1 in the WNT/β-catenin signaling pathway from the experiment described in **figure 8a**,**b**. GAPDH was used as a loading control. Data are combined from six PCLS samples pooled per donor. **a)** PCLS from non-PF donor tissue, and **b**) PCLS from PF donor tissue.

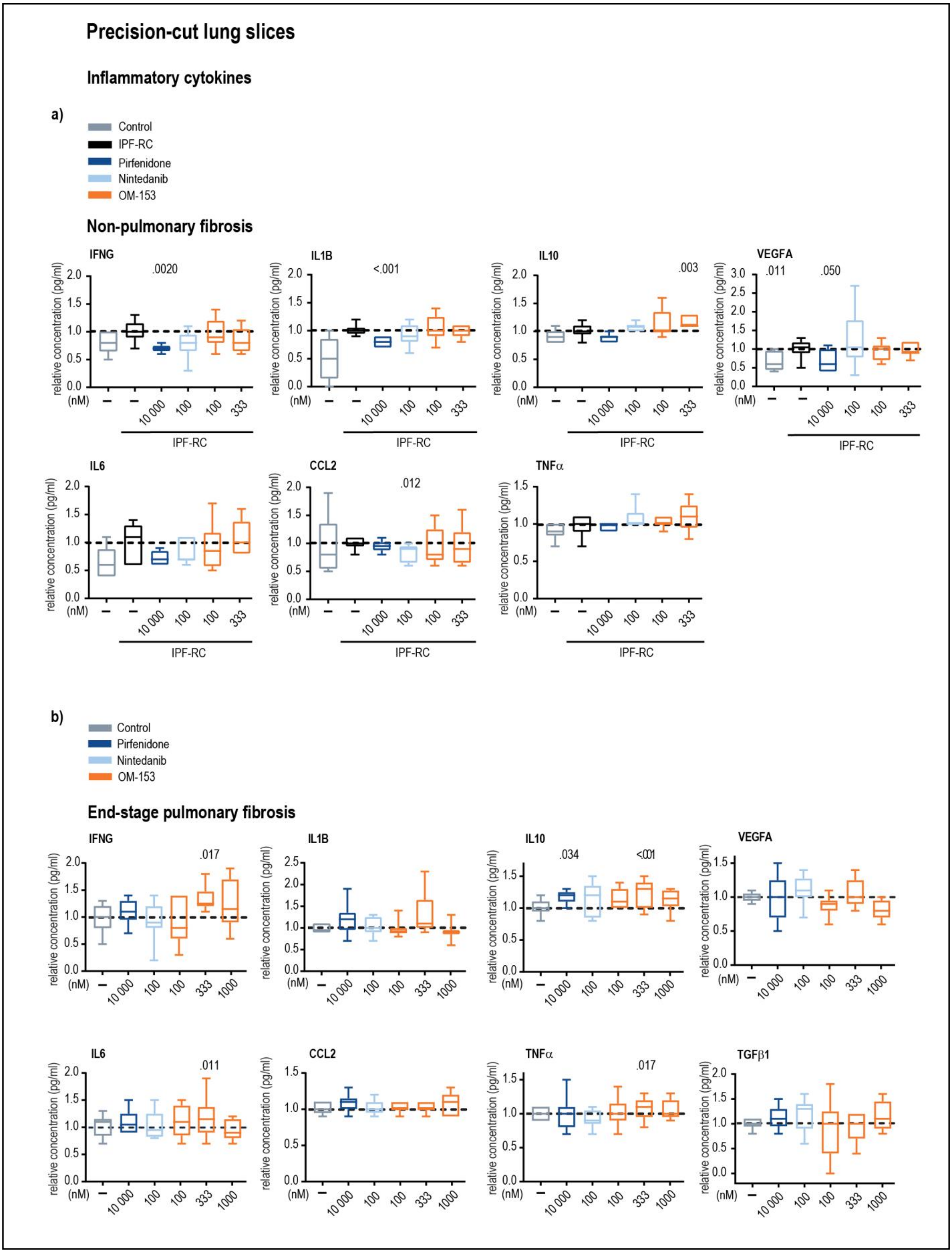
Precision-cut lung slices
Inflammatory cytokines
a)
Control
IPF-RC
Pirfenidone
Nintedanib
OM-153
Non-pulmonary fibrosis
IFNG
.0020
IL1B
<.001
IL10
.003
VEGFA
.011
.050
IL6
CCL2
.012
TNFα
relative concentration (pg/ml)
(nM)
10 000
100
100
333
IPF-RC
b)
Control
Pirfenidone
Nintedanib
OM-153
End-stage pulmonary fibrosis
IFNG
.017
IL1B
IL10
.034
<.001
VEGFA
IL6
.011
CCL2
TNFα
.017
TGFβ1
relative concentration (pg/ml)
(nM)
10 000
100
100
333
1000

**Supplementary Figure S11. OM-153 does not reduce inflammatory cytokines in *ex vivo* human lung fibrosis models**

**a**, **b)** Absolute concentrations (pg/mL) of inflammatory cytokines measured in culture supernatants from the experiment described in **figure 7**. Boxplots show median, first and third quartiles, and whiskers (min–max). Data are combined from three donors, each with six PCLS samples, pooled in pairs, with three replicates. Stippled lines represent mean IPF-RC or control values, respectively. *P*-values indicate two-tailed *t*-tests versus the respective controls. **a)** PCLS from non-PF donor tissue, and **b**) PCLS from PF donor tissue.

**Supplementary Table 1. OM-153 suppresses IPF-RC-induced profibrotic transcriptional programs in normal human lung fibroblasts**

Excel file containing RNA sequencing and IPA results, related to the analysis described in **figure 2**.

**Sheets 1–4)** DEGs identified from RNA-seq comparisons between control, IPF-RC, OM-153, nintedanib, and combination-treated samples, showing gene symbol, $\log_2$ fold change, and *P*-value.

**Sheets 5–7)** Predicted upstream regulators (UPSRs) from IPA for the same comparisons, showing upstream regulator, expression log ratio, activation z-score, and *P*-value of overlap. UPSRs were selected based on *P*-value < 0.05 and absolute activation z-score > 2.

**Sheets 8–10)** Canonical pathways (CPs) enriched by IPA, listing pathway name, activation z-score, and *P*-value. CPs were selected based on *P*-value < 0.05 and absolute z-score > 2.

**Sheets 11–13)** Gene clusters used for heatmap (HM) visualization, listing gene symbol and expression levels across conditions in $\log_2$ values.

**Sheets 14–17)** IPA output data for UPSRs and CPs generated using a combined input gene list from IPF-RC vs control and IPF-RC + OM-153 vs IPF-RC comparisons, corresponding to the analysis described in **figure 2e**. Data were selected using *P*-value < 0.05 and absolute z-score > 2.
