## Supplementary materials and methods for "Tankyrase inhibition demonstrates anti-fibrotic effects in preclinical pulmonary fibrosis models"

### Human lung fibroblast assays

NHLFs (#CC-2512, LOT 0000548315, Lonza, Basel, Switzerland) were cultured in high-glucose Dulbecco's Modified Eagle's Medium (DMEM, #31966021, Life Technologies, Carlsbad, CA, USA) supplemented with 10% fetal bovine serum (FBS, #10270-106, Life Technologies, Carlsbad, CA, USA) and 1% penicillin/streptomycin (P/S, #P4333, Sigma-Aldrich, St. Louis, MO, USA). Cell cultures were maintained below passage 20 (approximately 10 weeks) and monitored for mycoplasma contamination upon thawing and monthly thereafter using the MycoAlert™ Mycoplasma Detection Kit (#LT07-318, Lonza, Basel, Switzerland). IPF-RC was prepared as a 1000× stock in 0.1% bovine serum albumin (BSA, #A1595, Sigma-Aldrich, St. Louis, MO, USA). For *in vitro* experiments, NHLFs cultured in starvation medium (DMEM with 0.1% FBS and 1 % P/S) were stimulated with IPF-RC (1×) [1], TGFβ (0.3 ng/mL), or control (0.1% BSA) and treated with various concentration of OM-153 (Symeres, Nijmegen, Netherlands) or nintedanib (#656247-17-5; Kemprotec Ltd., Cumbria, UK) for 72 hours. For real-time qRT-PCR, RNA sequencing, and immunoblot analyses, NHLFs were seeded in 6-well plates (#140675, Nunc™, Roskilde, Denmark) at a density of $3 \times 10^5$ cells/well.  Immunoblotting, RNA isolation, and real-time qRT-PCR were performed as previously described [2]. The following additional TaqMan probes were used: *COL1A1* (Hs00164004_m1), *COL3A1* (Hs00943809_m1), *COL6A1* (Hs01095585_m1), *FN1* (Hs01549976_m1), and *ACTA2* (Hs00426835_g1), all from Thermo Fisher Scientific (Waltham, MA, USA). The Scar-in-a-Jar assay was performed as previously described [3] and the EarlyTox™ Live/Dead assay was conducted according to the manufacturer's instructions (#R8340, Molecular Devices, San Jose, USA). For the lung-on-a-chip assay, NHLFs were embedded at $10 \times 10^6$ cells/mL in fibrin and cultured as microtissues in uBeat® Stretch Platform for 7 days. Cells were initially maintained overnight in DMEM supplemented with 10% FBS and 2 mg/mL 6-aminocaproic acid (ACA, #A2504, Sigma-Aldrich, St. Louis, USA). The following day, medium was replaced with DMEM containing 0.1% FBS and 2 mg/mL ACA, alone or supplemented with IPF-RC (1×) or TGFβ (0.3 ng/mL). Treatments included OM-153 (33 nM) or nintedanib (100 nM). Medium was changed every other day with progressive ACA reduction (1.6 mg/mL on day 3, 1.2 mg/mL on day 5, and 1 mg/mL on day 7). Immunofluorescence staining was performed as previously described [4], , and mean fluorescence intensity was quantified within predefined ROIs using Otsu's algorithm thresholding.

**RNA sequencing and alignment**

After 72 hours of culture, NHLFs (Control, OM-153, IPF-RC, IPF-RC + nintedanib, and IPF-RC + OM-153) were detached using trypsin-EDTA (#T3924, Sigma-Aldrich, ST.Louis, MO, USA), and total RNA was isolated as previously described [2]. Samples at appropriate concentration and volume were sent to Novogene UK Cambridge Sequencing Center for mRNA sequencing with WBI-Quantification. Novogene performed RNA quality control, mRNA library preparation (poly A enrichment), and sequencing using the NovaSeq X Plus series PE150 (15 G raw data per sample). Raw FASTQ files were processed using the nf-core/rnaseq pipeline implemented in Nextflow (version 24.10.4). The reference genome was *Homo sapiens* GRCh38 (assembly: Homo_sapiens.GRCh38.dna_sm.primary_assembly.fa) with the corresponding GTF annotation (Homo_sapiens.GRCh38.113). The workflow included quality control, data filtering, mapping to the reference genome, and gene expression quantification.

**Bioinformatics**

Differential expression analysis was performed using DESeq2 in R (version 4.4.3) [5] comparing treatment conditions (n = 3 per group). Genes with a $p$-value < 0.01 and an absolute $\log_2$ fold change ≥ 0.3 were considered DEGs and included for downstream analysis. PCA, volcano plots, heatmaps, and GSEA were performed and visualized using R. PCA was used to explore variation between samples using variance-stabilizing data from the DESeq2 analysis, after batch adjustment with Limma [6, 7]. Volcano plots display the $\log_2$ fold change versus $-\log_{10}(p\text{-value})$ from the DESeq2 results. Genes with a $p$-value < 0.01 and an absolute $\log_2$ fold change ≥ 0.3 were highlighted. Heatmaps were created using z-scored $\log_2$-transformed expression values (n=3), with hierarchical gene clustering applied. Visualizations were produced using the ggplot2 package in R [8]. GSEA was conducted using gene sets from the Molecular Signatures Database (MSigDB) [9]. The selected gene sets included:
"GO_EXTRACELLULAR_MATRIX.v2024.1.Hs",
"GOBP_EXTRACELLULAR_MATRIX_DISASSEMBLY.v2024.1.Hs",
"HALLMARK_INFLAMMATORY_RESPONSE.v2024.1.Hs",
"REACTOME_COLLAGEN_FORMATION.v2024.1.Hs",
and "WP_LUNG_FIBROSIS.v2024.1.Hs".

Gene sets were restricted to those containing between 15 and 500 genes. For each DESeq2 contrast, genes were ranked by the Wald test statistic (stat). Enrichment results were visualized using a $q$-value threshold of 0.05, adjusted by the Benjamini–Hochberg method.

IPA® (QIAGEN Redwood City, CA, US) was performed using a combined list of 2,398 DEGs identified from the “Control vs IPF-RC" and “IPF-RC vs IPF-RC + OM-153" comparisons. These data were selected based on a *p*-value < 0.01 and a $\log_2$ fold change value ≥ ± 0.3. IPA analysis was conducted as previously described [10]. For the analysis, the following settings were applied: Expression Fold Change (Exp Fold Change), Relationships to consider (Direct and Indirect Relationships), Reference set (Corresponding data analysis), Interaction networks (35 molecules/network; 25 networks/analysis), Species (mammal: Human, mouse, rat). Molecule and Canonical Pathway subcategories were determined using all data types unless otherwise specified.

**Data and code availability**

The R code used for data analysis and visualization is publicly available on GitHub at https://github.com/MartinFStrand/TNKSiFibrosis.

**Mouse bleomycin-induced lung fibrosis model**

On day 1, male C57BL/6 mice (7-8 weeks old; 20230003012822, Shanghai Lingchang Biological Technology, Shanghai, China) received 1.5 mg/kg bleomycin (#600700, Nippon Kayaku, Tokyo, Japan) in saline via intratracheal (IT) instillation under isoflurane anesthesia. A sham group received IT saline only. On day 7, bleomycin-treated mice were randomly assigned to one vehicle control group and two treatment groups (n=10 per group). The treatment groups received either vehicle (5% dimethyl sulfoxide [DMSO, #D2650, Sigma-Aldrich, St. Louis, MO, USA], 50% polyethylene glycol 400 [PEG400, #81172, Sigma-Aldrich, St. Louis, MO, USA], and 45% saline, administered per-oral twice daily [PO-BID]), 30 mg/kg nintedanib (#T1777, Topscience, Shanghai, China, administered once daily [PO-QD]), or 10 mg/kg OM-153 (PO-BID, #PEZE25-084-1, Symeres, Nijmegen, Netherlands). The sham group was treated with vehicle only (n=5, PO-BID). Body weight and general health of all animals were monitored daily. Nine mice were euthanized for ethical reasons due to body weight loss exceeding 20%. Treatment continued until study termination on day 20. At endpoint, left lungs were harvested, fixed in 10% formalin (#HT501128, Sigma-Aldrich, St. Louis, MO, USA), embedded in paraffin (#39601095, Leica Biosystems, USA), and subjected to pathological analysis using Masson's Trichrome staining (#BA4079B) for fibrosis scoring [11] and H&E staining (#BA7621B) for injury scoring [12](both from Baso Diagnostics, Zhuhai, China). BALF was collected from the right lung for differential cell count as previously described [13] and for soluble collagen measurement using the Sircol™

soluble collagen assay (#S5000, Biocolor, Carrickfergus, UK). Immunoblotting, RNA isolation, and real-time qRT-PCR were performed as previously described [14]. The following antibodies were used for immunoblotting analysis: Anti-collagen type I (#sab4200678, Sigma-Aldrich, St. Louis, MO, USA), anti-collagen type III (#ab7778, Abcam, Cambridge, UK), and anti-collagen type IV (B-4, #sc-377143, Santa Cruz Biotechnology, Dallas, TX, USA).

**Human precision-cut lung slices**

PCLS were obtained from non-PF or end-stage PF donor tissue diagnosed with usual interstitial pneumonia (UIP) or non-specific interstitial pneumonia, as well as from tumor-free non-PF donor lung tissue from cancer patients [15]. All patients provided written informed consent, and the study was approved by the Ethics Committee of Hannover Medical School (Hannover, Germany, approval no. 2701–2015, renewed on 2015/04/22), in compliance with the “Code of Ethics of the World Medical Association”. Donor characteristics are summarized in **table 1**. PCLS were prepared according to previously established protocols [16-18]. PCLS from both PF and non-PF donors were treated with OM-153, nintedanib (#656247-17-5; Kemprotec Ltd., Cumbria, UK), or pirfenidone (#CT-PIRF, Chemietek, Indianapolis, USA) for 72 hours. Non-PF donor samples stimulated with 3× IPF-RC to induce fibrogenesis. Tissue viability after 72 hours was confirmed using lactate dehydrogenase (LDH)-based Cytotoxicity Detection Kit (#11644793001, Roche, Mannheim, Germany) [16]. Culture supernatants collected after 48 hours for enzyme-linked immunosorbent assay (ELISA) measurements, and tissues harvested at 72 hours was processed for RNA extraction [19] or protein extraction using RIPA buffer (#89900, Thermo Fisher Scientific, Waltham, MA, USA) and lysing matrix D tubes (#6913500, MP Biomedicals, Irvine, CA, USA) for immunoblotting.

### Table 1: Pathological findings in non-PF and PF donor lungs.

| Donor | Sex / Age (years) | Main histopathological findings | Comment / Interpretation |
|---|---|---|---|
| **Non-PF donor 1** | Female/77 | non-specific interstitial pneumonia. | - |
| **Non-PF donor 2** | Male/79 | non-specific interstitial pneumonia. | - |
| **Non-PF donor 2** | Male/52 | non-specific interstitial pneumonia. | - |
| **PF donor 1** | Female/58 | Advanced interstitial fibrosis with architectural distortion, bronchialization, myogenic metaplasia, and fibroblastic foci. Minor interstitial pneumonitis, chronic bronchitis, and mild pulmonary artery sclerosis. | Findings compatible with IPAF due to lympho-plasmacellular inflammation. No IgG4-related disease or malignancy. Stable compared with previous report. |
| **PF donor 2** | Male/61 | Advanced interstitial fibrosis with fibroblastic foci, chronic suppurative bronchitis/bronchiolitis, plasma cell– and eosinophil-rich pneumonitis, and low-grade pulmonary artery sclerosis. | Definite UIP pattern (ATS/ERS) consistent with IPF. Marked plasma cell and eosinophilic inflammation. No IgG4-related disease or malignancy. |
| **PF donor 3** | Male/60 | Advanced interstitial fibrosis with fibroblastic foci, chronic bronchitis/bronchiolitis, secretory retention, and mild pulmonary artery sclerosis. | Definite UIP pattern (ATS/ERS) consistent with clinically reported IPF. No malignancy. |

**ELISA and multiplex analyses**

For the scar-in-a.jar assay, the biomarkers quantifying collagen and fibronectin formation were measured in supernatant from the NHLFs using competitive ELISAs (Nordic Bioscience, Herlev, Danmark). The experiments were performed at Nordic Bioscience using NordicPRO-C3™ [20], NordicPRO-C6™ )[21], and NordicFBN-C™ [22] assays. Inflammatory cytokines in PLCS supernatants and whole mouse lung extracts were measured according to the manufacturer's instructions using the following kits from Thermo Fisher Scientific (Waltham, USA): Human ProcartaPlex Mix&Match 8-plex (PPX-08-MX2XAF4), Human ProcartaPlex TGFβ1 Simplex (EPX-01A-10249-901), ProcartaPlex Human Basic kit (EPX010-10420-901), Mouse ProcartaPlex Mix&Match 8-plex (PPX-08-MX323Z2), Mouse ProcartaPlex TGFβ1 Simplex (EPX01A-20608-901), ProcartaPlex Mouse Basic kit (EPX010-20440-901), Rat ProcartaPlex Mix&Match 8-plex (PPX-08-MX47XKY), Rat ProcartaPlex TGFβ1 Simplex (PX01A-30249-901), and ProcartaPlex Rat Basic kit (PX010-30420-901).
